# Structure of papain-like protease from SARS-CoV-2 and its complexes with non-covalent inhibitors

**DOI:** 10.1101/2020.08.06.240192

**Authors:** Jerzy Osipiuk, Saara-Anne Azizi, Steve Dvorkin, Michael Endres, Robert Jedrzejczak, Krysten A. Jones, Soowon Kang, Rahul S. Kathayat, Youngchang Kim, Vladislav G. Lisnyak, Samantha L. Maki, Vlad Nicolaescu, Cooper A. Taylor, Christine Tesar, Yu-An Zhang, Zhiyao Zhou, Glenn Randall, Karolina Michalska, Scott A. Snyder, Bryan C. Dickinson, Andrzej Joachimiak

**Affiliations:** Center for Structural Genomics of Infectious Diseases, Consortium for Advanced Science and Engineering, University of Chicago, Chicago, IL 60637, USA; Structural Biology Center, X-ray Science Division, Argonne National Laboratory, Argonne, IL 60439, USA; Department of Chemistry, University of Chicago, Chicago, IL, 60637, USA; Department of Microbiology, Ricketts Laboratory, University of Chicago, Chicago, IL, 60367, USA5; Department of Biochemistry and Molecular Biology, University of Chicago, Chicago, IL, 60637, USA

**Keywords:** Papain-like protease, PLpro, PLpro inhibitors, SARS-CoV-2, crystal structure, COVID-19

## Abstract

The number of new cases world-wide for the COVID-19 disease is increasing dramatically, while efforts to contain Severe Acute Respiratory Syndrome Coronavirus 2 is producing varied results in different countries. There are three key SARS-CoV-2 enzymes potentially targetable with antivirals: papain-like protease (PLpro), main protease (Mpro), and RNA-dependent RNA polymerase. Of these, PLpro is an especially attractive target because it plays an essential role in several viral replication processes, including cleavage and maturation of viral polyproteins, assembly of the replicase-transcriptase complex (RTC), and disruption of host viral response machinery to facilitate viral proliferation and replication. Moreover, this enzyme is conserved across different coronaviruses and promising inhibitors have already been discovered for its SARS-CoV variant. Here we report a substantive body of structural, biochemical, and virus replication studies that identify several inhibitors of the enzyme from SARS-CoV-2 in both wild-type and mutant forms. These efforts include the first structures of wild-type PLpro, the active site C111S mutant, and their complexes with inhibitors, determined at 1.60–2.70 Angstroms. This collection of structures provides fundamental molecular and mechanistic insight to PLpro, and critically, illustrates details for inhibitors recognition and interactions. All presented compounds inhibit the peptidase activity of PLpro *in vitro*, and some molecules block SARS-CoV-2 replication in cell culture assays. These collated findings will accelerate further structure-based drug design efforts targeting PLpro, with the ultimate goal of identifying high-affinity inhibitors of clinical value for SARS-CoV-2.

## INTRODUCTION

The Severe Acute Respiratory Syndrome Coronavirus 2 (SARS-CoV-2) is causing the COVID-19 pandemic. The virus belongs to the clade B of genus Betacoronavirus ^1^ and has large (+) sense ssRNA genome coding for 29 proteins. The 4 structural and 9-10 accessory proteins are translated from subgenomic RNAs produced from (–) sense ssRNA ^2^. To reach the replication stage, the CoV-2 genomic (+) sense ssRNA is used as mRNA to ultimately produce 15 non-structural proteins (Nsps) from two large polyproteins, Pp1a (4,405 amino acids) and Pp1ab (7,096 amino acids) ^3^. Pp1a is cleaved into the first 10 Nsps (Nsp11 is just a 7 residue peptide) and Pp1ab, which is made through a –1 ribosomal frame-shifting mechanism ^4^. The resulting Pp1ab contains all 15 Nsps ^5^. Therefore, proper polyprotein processing is essential for the release and maturation of the 15 Nsps and assembly into cytoplasmic, ER membrane-bound multicomponent replicase-transcriptase complex (RTC), which is responsible for directing the replication, transcription and maturation of the viral genome and subgenomic mRNAs ^6,7^.

There are two distinctive cysteine proteases encoded by the CoV-2 genome that are essential to the virus proliferation cycle ^6^: papain-like protease (PLpro, a domain within Nsp3, EC 3.4.22.46) and chymotrypsin-like main protease (3CLpro or Mpro, corresponding to Nsp5, EC 3.4.22.69). The main protease cuts 11 sites in Pp1a/Pp1ab with sequence consensus X-(L/F/M)-Q↓(G/A/S)-X ^7,8^ and PLpro cleaves 3 sites, with recognition sequence consensus “LXGG↓XX”, but is as indispensable as Mpro because its activity extends far beyond polyproteins cleavage.

PLpro is a domain of Nsp3 – a large multidomain protein that is an essential component of the RTC ^7,8^. The enzyme is located in Nsp3 between the SARS unique domain (SUD/HVR) and a nucleic acid-binding domain (NAB). It is highly conserved and found in all coronaviruses ^8^, often in two copies, denoted as PL1pro and PL2pro ^9,10^. In CoV-2, Nsp3 contains 1,945 residues (∼212 kDa) ^3^. This cysteine protease cleaves peptide bonds between Nsp1 and Nsp2 (LNGG↓AYTR), Nsp2 and Nsp3 (LKGG↓APTK), and Nsp3 and Nsp4 (LKGG↓KIVN) liberating three proteins: Nsp1, Nsp2 and Nsp3 ^10^. The LXGG motif found in Pp1a/Pp1ab corresponds to the P4–P1 substrate positions of cysteine proteases and is essential for recognition and cleavage by PLpro ^10,11^. Nsp1 is a 180 residue protein that interacts with 80S ribosome and inhibits host translation (https://doi.org/10.1101/2020.07.07.191676). Nsp2 is a 638 residue protein that was proposed to modulate host cell survival ^12^.

PLpro exhibits multiple proteolytic and other functions ^13^. In addition to processing Pp1a/Pp1ab, it was shown in SARS- and MERS-CoVs to have deubiquitinating activity, efficiently disassembling mono-, di-, and branched-polyubiquitin chains. It also has deISG15ylating (interferon-induced gene 15) activities. Both ubiquitin and ISG15 protein carry the PLpro recognition motif at their C-termini ^14,15^ suggesting that removal of these modifications from host cells interferes with the host response to viral infection ^10,16–19^. PLpro also inactivates TBK1, blocks NF-kappaB signaling, prevents translocation of IRF3 to the nucleus, inhibits the TLR7 signaling pathway, and induces Egr-1-dependent up-regulation of TGF-β1 ^18,20^. Further illustrating the complex and diverse functions of the protein, in some reports, various PLpro roles are decoupled from its proteolytic activity ^21,22^. Nevertheless, PLpro is a multifunctional protein having an essential role in processing of viral polyproteins, maturation, and assembly of the RTC, and it also may act on the host cell proteins by disrupting host viral response machinery to facilitate viral proliferation and replication. Due to the centrality of PLpro to viral replication, it is therefore an excellent candidate for therapeutic targeting.

Ongoing efforts to identify antivirals for CoV-2 to date have focused mainly on three Nsp proteins identified as the key drug targets from previous SARS- and MERS-CoV studies: Nsp3 PLpro, Nsp5 Mpro, and Nsp12 RNA-dependent RNA polymerase. Here we discuss the case for targeting SARS-CoV-2 Nsp3, PLpro. The enzyme is conserved in SARS-CoV, MERS-CoV, Swine Acute Diarrhea Syndrome (SADS) coronaviruses (Supplemental Fig. 1), and other viruses including Murine Hepatitis Virus, Avian Infectious Bronchitis Virus, and Transmissible Gastroenteritis Virus (TGEV); fortuitously, it has low sequence similarity to human enzymes. The sequence, structure, and functional conservation of PLpro suggests that therapeutics targeting SARS-CoV-2 PLpro may also be effective against related viruses with PLpro. In the past, this enzyme was structurally well characterized and currently there are over 40 structures of viral PLpro proteases in the Protein Data Bank ^23^, mainly from SARS-CoV, that can aid structure-based drug discovery. In fact, the past 15 years of studies with PLpro have led to the identification of a number of inhibitors that were specific for SARS-CoV PLpro, but did not inhibit the MERS-CoV enzyme ^7,24,25^. Unfortunately, these efforts have failed, thus far, to produce antivirals that can be useful for treatment of SARS-CoV-1 and −2 infections in humans.

Here we report eight crystal structures, including the first structure of wild-type PLpro, the active site cysteine mutant, and their complexes with known and novel inhibitors of SARS-CoV-2 PLpro, determined at 1.60–2.70 Angstroms. These data reveal the structural basis of the enzyme with fine molecular details, and illustrate specific ligand recognition and interactions. The presented compounds inhibit PLpro peptidase activity *in vitro*, and most importantly, several of the novel inhibitors also block SARS-CoV-2 replication in cell culture. Collectively, these findings provide critical insights for further structure-based drug design efforts against PLpro to enable the design of even higher affinity inhibitors and, ultimately, human therapeutics.

## Results and Discussion

The CoV-2 PLpro sequence is 83% identical and 90% similar to SARS-CoV-1 and 31% identical and 49% similar to MERS-CoV and even more distant to SADS PLpro (Supplemental Fig. 1). Between SARS- and MERS-CoV many substitutions are quite conservative, but there are some that may have significant impact on protein stability, dynamics, ligand binding, and catalytic properties. Examples include Thr75 (Leu/Val), Pro129 (Ala/Ile), Tyr172 (His/Thr), Lys200 (Thr/Gln), Lys274 (Thr/Val) and Cys284 (Arg/Arg), with equivalent residues shown in parentheses for SARS/MERS PLpro, respectively.

CoV-2 PLpro is a slightly basic, 315 residue protein with high content of cysteine residues (3.5%). In addition to catalytic Cys111, there are four cysteine residues coordinating important structural zinc ion and other six distributed throughout the protein structure. Similar to Mpro (submitted) ^26^, the active site cysteine seems much more reactive as evident by structures of covalent adducts reported in the PDB (PDB id: 6WX4 and 6WUU). CoV-2 Mpro-active cysteine has been shown to have different level of oxidation in crystals (PDB id: 6XKF, 6XKH and publication submitted). The potential sensitivity of PLpro Cys111 to oxidation presented a challenge for structure determination as the wild-type protein exhibited rather poor crystallization properties. The PLpro active site contains a canonical cysteine protease catalytic triad (Cys111, His272 and Asp286) ^17^, while Mpro has catalytic dyad (Cys145 and His41) ^26^, which may account for somewhat dissimilar chemical properties of the two enzymes. PLpro may have catalytic properties more common with other cysteine proteases with the generally accepted thiolate form of Cys111 acting as nucleophile, His272 serving as a general acid-base, and Asp286 promoting deprotonation of Cys111.

### Structure determination and structural comparisons

We have determined eight structures of PLpro from CoV-2, including two wild-type apo-protein structures, one at 2.70 Å (PDB id: 6W9C) and the other in different crystal packing at 1.79 Å (PDB id: 6WZU), both at 100 K, the apo-PLpro active site C111S mutant under cryogenic conditions 100 K at 1.60 Å (PDB id: 6WRH) and at 293 K at 2.50 Å (PDB id: 6XG3). The first structure (WILD-TYPE) was solved by molecular replacement using SARS PLpro model as a template. The subsequent structures were phased with the refined wild-type model. Structures were refined as described in Supplementary Materials and Methods and data and refinement statistics is shown in Supplementary Table 1. For high resolution structures, all residues are visible in the electron density maps, for 2.70 Å structure of wild-type enzyme three N-terminal residues are missing and for the 293 K 2.50 Å mutant structure two N-terminal and one C-terminal residues are missing. Of significance, the electron density map for residues around the active site is excellent. These structures are virtually identical with the largest difference being between the lowest (2.70 Å) and highest resolution (1.60 Å) structures (RMSD 0.72 Å). They differ the most in the zinc-binding region and in the Gly266 – Gly271 loop containing Tyr268 and Gln269. The high resolution wild-type and C111S mutant structures show RMSD 0.10 Å, and 293 K and 100 K mutant structures show RMSD 0.27 Å. In the high resolution structures, in addition to structural zinc ion, there are two chloride and two phosphate ions bound. In the highest resolution structure we modeled 381 water molecules.

Structures of Nsp3 PLpro were reported previously for SARS-, MERS-CoV, and other viruses ^27^. The CoV-2 PLpro structure has “thumb–palm–fingers” architecture described before (Fig. 1). This arrangement is similar to the ubiquitin specific proteases (USPs), one of the five distinct deubiquitinating enzyme (DUB) families, despite low sequence identities (∼10%) ^10,27^. Briefly, the protein has two distinct domains: the small N-terminal ubiquitin-like (Ubl) domain and the “thumb–palm–fingers” catalytic domain (Fig. 1). The Ubl domain consists of residues 1 to 60 with five β-strands, one α-helix, and one 3_10_-helix. In CoV-2 PLpro, chloride ion binds to a small loop formed by residues Thr9 – Ile14 at the interface with the catalytic domain. In different structures, Ub1 shows some conformational flexibility, though the specific function of this domain is not well understood.

**Figure 1.**
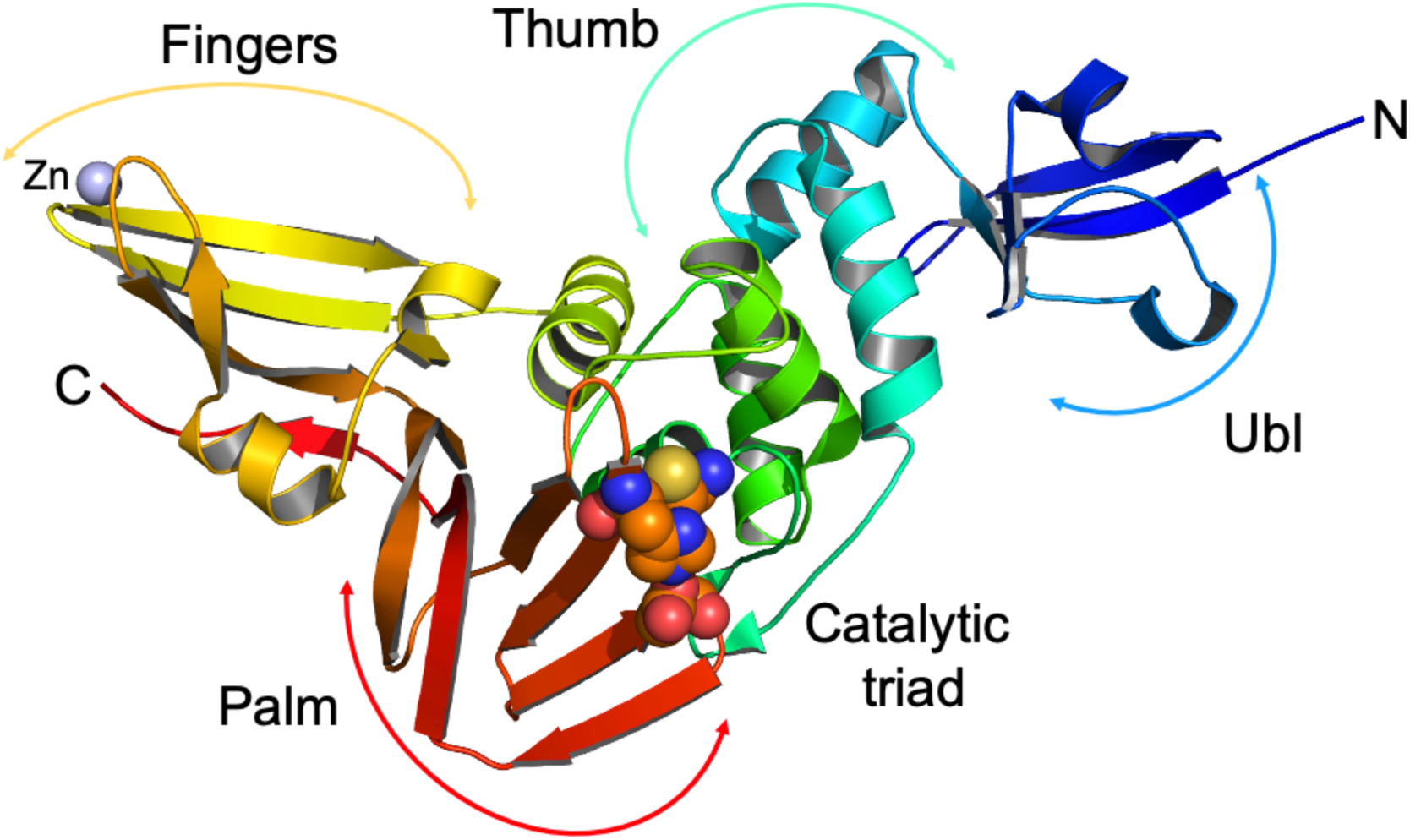
Structure of PLpro from SARS-CoV-2 showing secondary structure, domains and subdomains with active site residues Cys111/His272/Asp286 represented as spheres, zinc ion is in blue.

The larger catalytic domain is an extended right-hand scaffold with three characteristic subdomains. A thumb is comprised of six α-helices and a small β-hairpin. The fingers subdomain is the most complex; it is made of six β-strands and two α-helices and includes a zinc binding site. This structural zinc ion is coordinated by four cysteines located on two loops (Cys189, 192, 224, and 226) of two β-hairpins. Zinc binding is essential for structural integrity and protease activity ^28^, but the conformation of this region varies most between different PLpro structures. The palm subdomain is comprised of six β-strands (Fig. 1) with the catalytic residues Cys111, His272, and Asp286 located at the interface between the thumb and palm subdomains (Fig. 1). An important mobile β-turn/loop (Gly266 – Gly271) is adjacent to the active site that closes upon substrate and/or inhibitor binding. In the high resolution structure of PLpro C111S mutant, there is a phosphate ion bound to the active site at the N-terminus of helix α4 (contributing Cys111) that is coordinated by Trp106, Asn109, and His272. This site provides a good environment to stabilize C-terminal carboxylate group of the peptide cleavage product. In this structure, there is another phosphate ion bound to a thumb subdomain and is coordinated by His73 and His170. This subdomain also binds a second chloride ion near Arg140. In the structures with inhibitors there are additional zinc and chloride ions bound, including one in the active site that is coordinated by the active site Cys111.

We compared our structures with the high resolution crystal structures of SARS and MERS PLpro. The SARS PLpro Cys112Ser mutant in complex with ubiquitin (PDB id: 4M0W) shows RMSD 0.53 Å with our highest resolution structure of CoV-2 PLpro C111S mutant (PDB id: 6WRH). The largest differences are observed, in our lower resolution wild-type structure, in zinc binding region, consistent with this region being the most flexible in the PLpro structures. A comparison of MERS PLpro high resolution structure (PDB id: 4RNA) with CoV-2 PLpro C111S mutant shows much bigger differences (RMSD 1.82 Å), with the largest structural shifts occurring again in the structural zinc-binding region and also in the N-terminal Ubl domain. The PLpro core shows analogous differences with RMSD 1.72 Å (PDB id: 5KO3). In compared structures, as expected, side chains of many surface residues show different conformations. Nevertheless, the arrangement of the catalytic site is very similar in SARS-CoV-2 and SARS-CoV, suggesting that at least some inhibitors may display cross activity between these proteases. The MERS PLpro active site region differs quite significantly from SARS PLpro enzymes and at least some SARS-specific inhibitors may not cross react.

### Enzyme activity assays of synthetic compounds

Several naphthalene-based compounds were synthesized (Supplemental Fig. 2) and tested for inhibition of SARS-CoV-2 PLpro activity (Supplemental Fig. 3). One of these compounds (**1**/ GRL0617) was identified previously as a specific SARS-CoV PLpro inhibitor and showed good potency and low cytotoxicity in SARS-CoV-infected Vero E6 cells ^29^. We present in this manuscript results of biochemical, whole cell, and high resolution crystallographic studies of seven compounds, six possessing the methyl-*N*-[(1*R*)-1-naphthalen-1-ylethyl]benzamide scaffold and one being a simplified analog of our own design. All these compounds inhibit SARS-CoV-2 PLpro protease and they are designated as follows: **1** is 5-amino-2-methyl-*N*-[(1*R*)-1-naphthalen-1-ylethyl]benzamide (GRL0617), **2** is 5-carbamylurea-2-methyl-*N*-[(1*R*)-1-naphthalen-1-ylethyl]benzamide, **3** is 5-acrylamide-2-methyl-*N*-[(1*R*)-1-naphthalen-1-ylethyl]benzamide, **4** is 3-amino-*N*-(naphthalene-1-yl)-5-trifluoromethyl)benzamide, **5** is 5-(butylcarbamoylamino)-2-methyl-*N*-[(1*R*)-1-naphthalen-1-ylethyl]benzamide, **6** is 5-(((4-nitrophenoxy)carbonyl)amino)-2-methyl-*N*-[(1*R*)-1-naphthalen-1-ylethyl]benzamide, and **7** is 5-pentanoylamino-2-methyl-*N*-[(1*R*)-1-naphthalen-1-ylethyl]benzamide (Fig. 2, Supplemental Fig. 2A).

**Figure 2.**
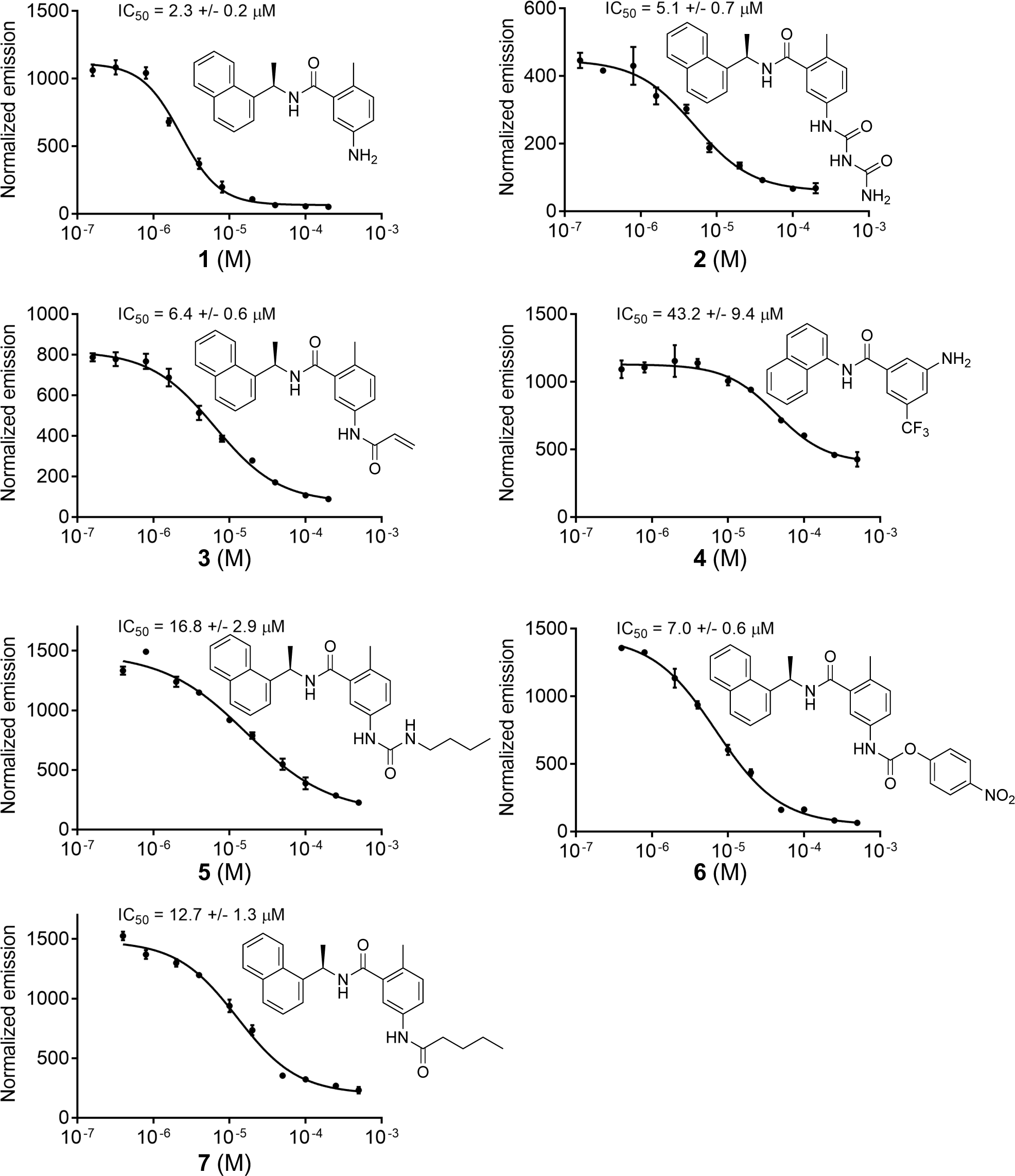
Biochemical activity assays for compounds **1** – **7** using substrate shown in supplemental Fig. 3.

We have developed an *in vitro* biochemical assay for PLpro using expressed protein and a pro-fluorescent peptide substrate, CV-2, designed based on the LKGG recognition motif of PLpro (Supplemental Figs. 2B and 3). CV-2 generates fluorescent signal in response the protease activity of PLpro, and critically, is unresponsive in the C111S variant (Supplementary Fig. 3). In this assay all seven compounds act as non-covalent inhibitors of the enzyme. Given that compound **1/**GRL0617 is known to inhibit SARS-CoV PLpro with an IC_50_ value of 0.6 μM ^29^, we expected it would likely inhibit the SARS-CoV-2 enzyme and indeed it does so, with an IC_50_ value of 2.3 μM in our assay conditions (Fig. 2). Compounds **2**, **3**, **5**, **6**, and **7** are further amine-functionalized derivatives of **1/**GRL0617, while **4** is a simplified variant of **1** without a chirality center (Supplemental Fig. 2A); despite their structural differences, we found that all inhibit PLpro to varying degrees (IC_50_ = 5.1–32.8 μM, Fig. 2). Given this suite of molecules that function as PLpro inhibitors *in vitro*, we next sought to test whether these molecules are also capable of inhibiting viral replication in live cells.

### Whole cell virus replication assays

We next performed CoV-2 virus replication assays using Vero E6 cells and measuring SARS CoV-2 replication. Surprisingly, not all compounds functioned in this assay, and their relative abilities to inhibit viral replication did not necessarily correlate directly with *in vitro* inhibition parameters toward PLpro. Compounds **1**, **4**, **5**, **6**, and **7** all proved capable of affecting the viability of cells and inhibiting virus replication. For compounds **1, 4,** and **7** their EC_50_ values range from 1.5 to 8.0 μM (Fig. 3A and 3B, and Supplemental Fig. 4). For example, PLpro inhibition by compound **1/**GRL0617 is 14-times better than **4**, but inhibition of virus replication is only two times higher. Moreover, compounds **2** and **3** are quite good inhibitors (IC_50_ values of 5.1 and 6.4 μM, respectively), but failed in the viral replication assay. Compound **5** was the weakest inhibitor *in vitro* (IC_50_ values of 32.8 μM), but was one of the best performers in the live viral replication assay (EC_50_ 2.5 μM).

**Figure 3.**
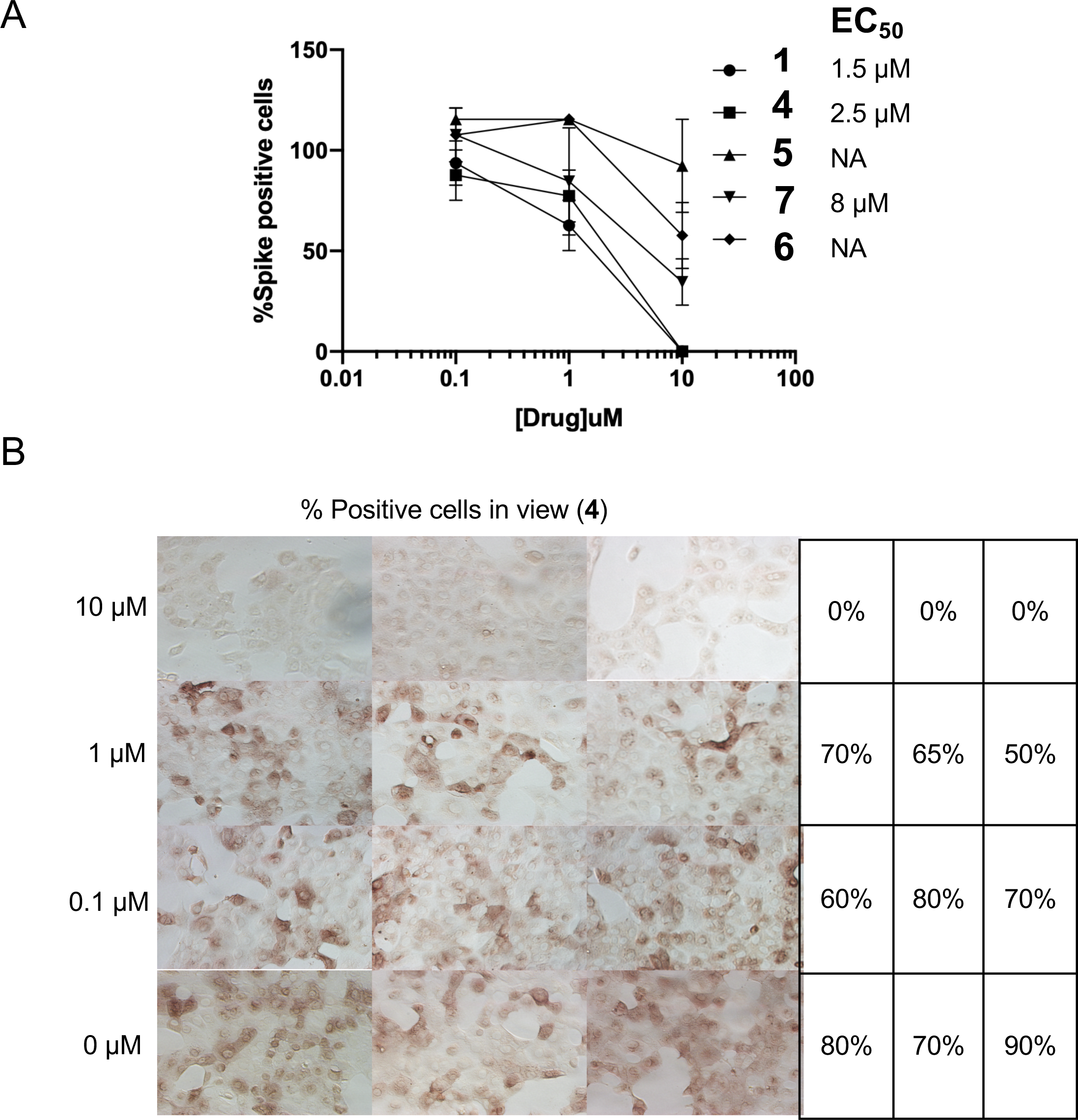
Virus inhibition in whole cell assay. A) Virus replication activity assays for compounds **1**, **4**, **5**, **6** and **7**. B) whole cell assay for compound **4**.

Differences in cell permeability and solubility could account for the disconnects between the *in vitro* biochemical assay data and viral replication data, but given the high degree of structural similarity between these molecules, these data indicate that further optimization is possible, especially in the case of compound **5**, which is a relatively weak binder but solid inhibitor of the virus. More broadly, all of the compounds are promising and may need only small modification(s) in order to serve as preclinical lead compounds. To enable structure-based improvements of these molecules, we next aimed to get ligand-bound structures of as many of these lead compounds as possible. Based on these results, we believe it is critical to combine the *in vitro* biochemical assays to triage compounds with live viral replication assays.

### Crystal structures with bound compounds 1, 2 and 3

We were able to determine crystal structures of SARS-CoV-2 PLpro with non-covalent inhibitors **1**, **2**, and **3** (Supplemental Table 1) three using C111S mutant and one wild-type enzyme (PDB ids: 7JIR, 7JIT, 7JIV, 7JIW). The structure with compound **3** was solved in both wild-type and mutant forms. The electron density for the ligand, protein, solvent, and bound acetate ion is excellent (Fig. 4). All three compounds bind to the same site as observed previously for GRL0617 in complex with SARS-CoV PLpro (PDB id: 3E9S) ^29^ and now determined for **1** in complex with SARS-CoV-2 PLpro (Fig. 4A). The structure of **2** bound to the PLpro C111S mutant was determined at 1.95 Å (PDB id: 7JIT), the highest resolution for all complexes to date, and this structure will be used here as a reference (Fig. 4B and Supplemental Fig. 5A). Compound **2** binds to the groove on the surface of PLpro near the active site, ∼8 Å apart from Cys/Ser111, overlapping with S4/S3 protein subsites (corresponding the substrate positions P4/P3), that are critical for recognition of the leucine residue in the LXGG motif (Fig. 4E). The ubiquitin peptide-binding site is a solvent-exposed groove: wide at S4 site, solvent exposed at P3 site, and very narrow at P2 and P1 sites (Fig. 4E). Because of the high resolution achieved, we observe extensive interactions between compound **2** and the protein involving direct as well as water-mediated hydrogen bonds and van der Waals contacts, noting some additional interactions provided by the carbamylurea moiety in **2**. As compared with unliganded protein, the main chain and side chains of several residues significantly adjust to accommodate the ligand (Arg166, Glu167, Tyr268, Gln269, Fig. 4B and Supplemental Fig. 5A). Direct hydrogen bonds are found between Glu167 and Tyr268 and two nitrogen and oxygen atoms of the carbamylurea moiety, respectively (Fig. 4B). As compared with **1**/GRL0617 compound **2** makes all the same interactions and provides additional four hydrogen bonds. Intriguingly, however, the IC_50_ value for **1** is ∼2 times lower than for **2**, despite **2** making more interactions with the protein. Asp164 hydrogen bonds with another amino group in the linker between two aromatic rings and main-chain amino group of Gln269 hydrogen bonds to oxygen atom of that linker. Lys157 makes water-mediated hydrogen bond to the carbamylurea moiety. This water molecule also coordinates Glu167. Interestingly, this residue (equivalent of Glu168 in SARS PLpro) seems to play an important role in Ub1 core recognition, and mutations can cause a significant loss of DUB activity.

**Figure 4.**
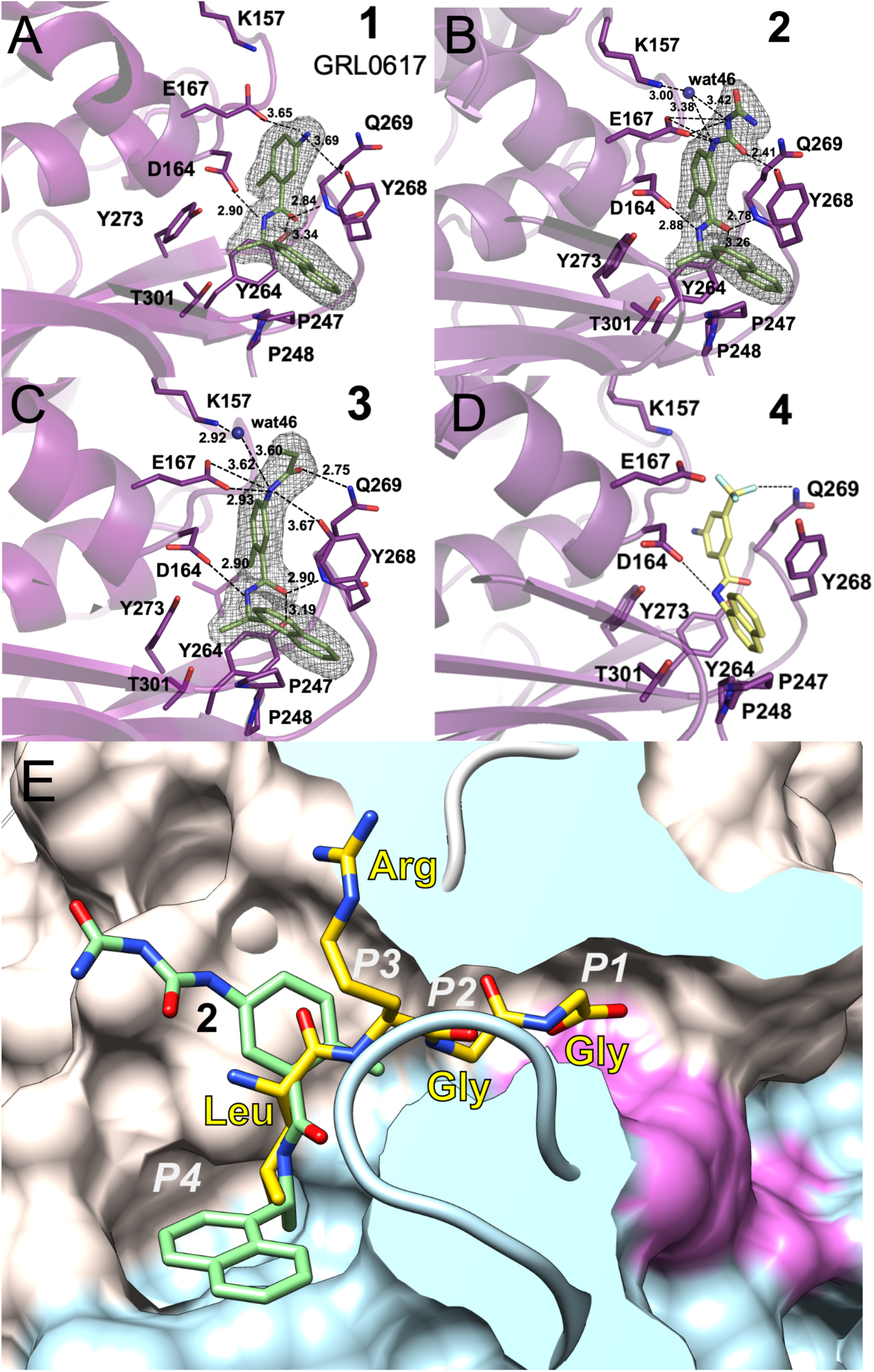
Ligands (green sticks) binding to SARS-CoV-2 PLpro (in magenta). A) Compound **1** binding to PLpro. B) Compound **2** binding to PLpro (in magenta). C) Compound **3** binding to PLpro (in magenta). D) Model of compound **4** (yellow sticks) binding to PLpro. Dashed lines show hydrogen bonds, water molecules are shown as blue spheres. In A, B and C the 2F_o_-mF_c_ electron density maps are shown as a grey mesh, contoured at 1.2 σ. E. Compound **2** (green sticks) binds to a groove on the surface of PLpro protein (surface of palm subdomain is in white and thumb subdomain is in light blue) with the active site catalytic triad surface is shown in red in the end of a slender tunnel. Peptide LRGG from ubiquitin structure in complex with SARS PLpro (PDB id: 4MOW) is shown in yellow and peptide positions corresponding P1 – P4 sites are marked in white.

The aromatic rings of **2** make several hydrophobic interactions, specifically with Pro248, Tyr268, aliphatic regions of the side chain of Gln269, and Asp164. Curiously, there is also an acetate ion packing in between **2** and protein residues. This ion is coordinated by Arg166 and Glu167. The ligand binding site offers a number of opportunities to improve ligand affinity (water and acetate binding site) and potential linking to the active site for covalent attachment, many of which were not explicitly indicated by previous work with SARS.

The structures of inhibitor **3** were determined in both forms: wild-type at 2.30 Å (PDB id: 7JIW) and C111S mutant at 2.05 Å (PDB id: 7JIV) (Fig. 4C). The structures of protein, the pose, and overall interactions of the inhibitor are the same in both variants. Compound **3** is a derivative of **1**, in which its amino moiety is functionalized to become an acrylamide. Structural comparison shows that only some interactions are preserved – all hydrophobic interactions and one hydrogen bond. The acrylamide moiety in **3** provides two additional hydrogen bonds as compared to **1**. Thus far, we have not been able to grow crystals of SARS-CoV-2 PLpro with compound **4**. However, we were able to model the interaction of **4** with the protein (Fig. 4D), where it appears that the trifluoromethyl moiety is able to interact with the amide group of Gln269. Given that several other analogs of **4** (structures not shown here) are inactive in biochemical studies, this interaction might be significant.

The inhibitory effect of compounds **1**, **2**, **3**, and **4** can be easily rationalized as they anchor to the site. Although somewhat away from the catalytic triad, their binding still interferes with the recognition of peptide motif LXGG. Comparison with the high resolution structure of PLpro with ubiquitin (PDB id: 4M0W) shows that the inhibitor linker region connecting naphthalene and benzene rings overlaps with the leucine residue of ubiquitin C-terminal sequence bound to the S4 subsite (Fig. 4E and supplemental Fig. 5B). The S4 site is where specificity of the LXGG peptide is determined as it recognizes leucine side chain by fitting it into hydrophobic pocket formed by Pro248, Tyr264, Tyr272 and Thr301. These residues are conserved in SARS CoV-1 and CoV-2 PLpro, but only Tyr272 and Thr301 are conserved in SARS and MERS PLpro (Supplemental Fig. 1). Therefore, ligands that bind to hydrophobic S4 site and make hydrophilic interactions with PLpro surface residues may be good candidates for inhibitors of the PLpro enzyme. The S3 site can accept any residue because it is solvent exposed, but would prefer hydrophilic side chain (Arg, Lys, Asn). The peptide then follows the path to the active site that becomes narrower and can accept only two glycine residues in P1 and P2. The P1’ again is on the protein surface and can accept any residue. The interesting requirement is that the peptide binding to S1 – S4 sites must be in extended/linear conformation, placing noteworthy constrains on designing inhibitors targeting the active site.

## Conclusions

In summary in this report we have presented a substantial high resolution structural, biochemical, and virus replication studies of PLpro cysteine protease from SARS-CoV-2 and describe seven compounds that inhibit enzyme in *in vitro* biochemical assay based on the cleavage of the LXGG recognition peptide, a subset of these inhibit virus replication. We have determined apo-structures of wild-type enzyme and inactive mutant in which single sulfur atom was replaced by oxygen (C111S). The apo and mutant structures were determined at high resolution 1.79 and 1.60 Angstroms providing detailed and accurate three-dimensional models of the enzyme. The mutant structures were determined under both 100K and 293K temperatures providing information about protein flexibility. All four apo-PLpro structures showed significant structural conservation, including catalytic triad and ordered solvent molecules. The protein surface provides rich chemical environment and is capable of binding a variety of ions including conserved structural zinc ion, few non-structural zinc ions, several phosphate, chloride and acetate anions. The structures of complexes with inhibitors **1**, **2**, and **3** were determined at 1.95–2.30 Angstroms, including compound **3** in both wild-type and mutant forms. All three ligands bind to the same site in the enzyme located 8 - 10 Å away from the catalytic cysteine and it is expected that all seven synthetic compounds bind in a very similar manner. Based on this assumption we have modelled pose of compound **4** in the structure. Considerable conformational adjustments are observed for the side chains of residues involved in ligand binding. These inhibitors bind to protease S4/S3 sites. The S4 site is where specificity of the LXGG sequence recognition motif is determined as it recognizes leucine side chain by fitting it into hydrophobic pocket. This site is only partly conserved between SARS CoV and MERS CoV enzymes explaining lack of cross reactivity of compound **1** reported previously. Binding inhibitors to this site would block peptide recognition. The PLpro peptide binding site narrows significantly as it approaches catalytic triad explaining why only glycine residues are accepted at the C-terminus of the recognition motif. These compounds would not only prevent virus polyproteins processing but also cleavage of host proteins modifications with ubiquitin and ISG15, therefore inhibit several PLpro functions. Five out of seven inhibitors of PLpro block virus replication in whole cell assay. Interestingly, their relative abilities to inhibit viral replication do not directly correlate with *in vitro* inhibition of PLpro, suggesting that other factors are important. Nevertheless our studies showed potential S4/S3 site binders to serve as scaffolds for effective inhibitors of SARS CoV-2 coronavirus. Our collection of structures provides fundamental molecular and mechanistic insight into PLpro structure and it illustrates details ligand recognition and interactions. These collated findings will accelerate further structure-based drug design efforts targeting PLpro, with the ultimate goal of identifying high-affinity inhibitors of clinical value for SARS-CoV-2.

## Acknowledgements

We thank the members of the SBC at Argonne National Laboratory, especially Darren Sherrell and Alex Lavens for their help with setting beamline and data collection at beamline 19-ID. Funding for this project was provided in part by federal funds from the National Institute of Allergy and Infectious Diseases, National Institutes of Health, Department of Health and Human Services, under Contract HHSN272201700060C (to AJ) and by the DOE Office of Science through the National Virtual Biotechnology Laboratory, a consortium of DOE national laboratories focused on response to COVID-19, with funding provided by the Coronavirus CARES Act (to AJ). The use of SBC beamlines at the Advanced Photon Source is supported by the U.S. Department of Energy (DOE) Office of Science and operated for the DOE Office of Science by Argonne National Laboratory under Contract No. DE-AC02-06CH11357. Funding for the synthesis and biochemical studies was provided by a “BIG” Award from the University of Chicago, the University of Chicago Women’s Board, the National Institutes of Health (TM GM08720, Predoctoral Training Program in Chemistry and Biology, graduate fellowship to CAT), the National Institute of General Medical Sciences (R35 GM119840 to BCD), and start-up funds from the University of Chicago (SAS).

## Author Contributions

AJ initiated the project, ME and RJ cloned and expressed wild-type and mutant proteins, RJ purified the first batch of protein and obtained the first wild-type PLpro crystal. CT purified proteins and crystallized proteins and complexes, while JO collected diffraction data, determined, refined and analyzed structures. YK contributed to structure refinement and analysis of structural data. VGL, SLM, SAS, CAT, YZ, and ZZ designed and synthesized compounds **1**–**7**. SAA and RSK synthesized CV-2. KJ performed all *in vitro* biochemical assays. GR, VN and SD performed virus assay and analyzed data. Finally, AJ, BCD, and SAS conceived of and directed the research as well as wrote the manuscript, while KM analyzed structural data and also wrote portions of the manuscript.

---

**Supplemental Table 1.** Data processing and refinement statistics.

|  | **PLpro WT**<br>**100K**<br>**2.70 Å** | **PLpro WT**<br>**100K**<br>**1.79 Å** | **PLpro C111S**<br>**mutant 100K**<br>**1.60 Å** | **PLpro C111S**<br>**mutant RT**<br>**2.50 Å** | **PLpro C111S**<br>**mutant – 1**<br>**100K, 2.09 Å** | **PLpro C111S**<br>**mutant – 2**<br>**100K , 1.95 Å** | **PLpro C111S**<br>**mutant – 3**<br>**100K, 2.05 Å** | **PLpro**<br>**WT – 3**<br>**100K, 2.30 Å** |
| --- | --- | --- | --- | --- | --- | --- | --- | --- |
| Wavelength (Å) | 0.9792 | 0.9792 | 0.9792 | 0.9792 | 0.9792 | 0.9792 | 0.9792 | 0.9792 |
| Resolution<br>range (Å) | 43.7-2.70<br>(2.77-2.70) | 48.8-1.79<br>(1.840–1.793) | 49.0-1.60<br>(1.641-1.600) | 49.3-2.50<br>(2.54-2.50) | 49.7-2.09<br>(2.13-2.09) | 49.7-1.95<br>(1.98-195) | 45.4-2.05<br>(2.09-2.05) | 45.5-2.3<br>(2.34-2.30) |
| Space group | C2 | P3<sub>2</sub>21 | P3<sub>2</sub>21 | P3<sub>2</sub>21 | I4<sub>1</sub>22 | I4<sub>1</sub>22 | I4<sub>1</sub>22 | I4<sub>1</sub>22 |
| Unit cell (Å, °) | $a = 191, b = 110, c = 64, \beta = 96$ | $a = 82, c = 134$ | $a = 82, c = 135$ | $a = 82, c = 135$ | $a = 114, c = 220$ | $a = 114, c = 219$ | $a = 114, c = 220$ | $a = 114, c = 220$ |
| Unique<br>reflections | 20799 (679) | 49598 (2456) | 69708 (3105) | 19357 (942) | 42950 (2097) | 52219 (2,368) | 45458 (2245) | 32308 (1576) |
| Redundancy | 2.5 (1.9) | 14.2 (12.4) | 13.7 (8.3) | 9.8 (8.9) | 12.6 (11.5) | 16.6 (9.5) | 21.0 (15.2) | 20.9 (14.6) |
| Completeness<br>(%) | 57.3 (38.4) | 100 (100) | 99.1 (89.5) | 100 (100) | 99.8 (99.3) | 99.5 (91.6) | 99.9 (99.6) | 99.9 (99.0) |
| Mean I/sigma(I) | 6.1 (1.14) | 27.2 (1.72) | 34.8 (1.71) | 17.2 (1.4) | 21.0 (1.07) | 35.2 (1.08) | 42.4 (1.03) | 44.3 (0.97) |
| Wilson B-factor<br>(Å<sup>2</sup>) | 46.9 | 23.0 | 33.6 | 54.4 | 51.8 | 36.3 | 49.2 | 66.3 |
| R-merge<sup>b</sup> | 0.140 (0.585) | 0.137 (1.78) | 0.095 (0.871) | 0.167 (1.61) | 0.145 (1.94) | 0.097 (1.23) | 0.101 (1.71) | 0.095 (1.73) |
| CC1/2<sup>c</sup> | 1.01 (0.566) | 0.991 (0.632) | 0.996 (0.741) | 0.987 (0.514) | 0.995 (0.512) | 1.00 (0.559) | 1.00 (0.566) | 1.01 (0.605) |
| **Refinement** |  |  |  |  |  |  |  |  |
| R-value (all)<br>(%) | 23.89 | 16.07 | 12.55 | 15.4 | 18.65 | 17.6 | 18.8 | 21.4 |
| R<sub>work</sub> value (%) | 23.51 (33.3) | 16.00 (25.0) | 12.34 (22.3) | 15.1 (29.3) | 18.58 (32.0) | 17.5 (30.9) | 18.8 (34.1) | 21.2 (38.9) |
| Free R-value<br>(%) | 30.88 (39.2) | 17.41 (26.4) | 16.43 (26.5) | 19.3 (29.6) | 29.98 (35.0) | 19.0 (30.5) | 20.1 (34.0) | 23.9 (39.5) |
| **Rms deviations** |  |  |  |  |  |  |  |  |
| Bond length (Å) | 0.004 | 0.010 | 0.009 | 0.010 | 0.010 | 0.011 | 0.010 | 0.006 |
| Angle (°) | 1.32 | 1.49 | 1.43 | 1.58 | 1.62 | 1.65 | 1.68 | 1.49 |
| Chiral (Å) | 0.044 | 0.070 | 0.071 | 0.066 | 0.069 | 0.072 | 0.073 | 0.059 |

| No. of atoms and mean B-factor (Å<sup>2</sup>) |  |  |  |  |  |  |  |  |
| --- | --- | --- | --- | --- | --- | --- | --- | --- |
| Protein | 7392 | 2575 | 2599 | 2539 | 2495 | 2561 | 2496 | 2426 |
| Ligand | - | - | - | - | 23 | 29 | 27 | 27 |
| Zn | 3 | 1 | 1 | 1 | 4 | 4 | 4 | 4 |
| Cl | 1 | 2 | 2 | 1 | 4 | 3 | 3 | 4 |
| Acetate | - | - | - | - | 4 | 8 | 4 | - |
| MES | - | - | - | - | 12 | 12 | 12 | 12 |
| PO<sub>4</sub> | - | 10 | 15 | 5 | - | - | - | - |
| Glycerol | - | 12 | 12 | - | - | - | - | - |
| Water | 1 | 223 | 380 | 41 | 127 | 210 | 135 | 32 |
| All atoms | 59.2 | 39.3 | 31.4 | 59.7 | 67.7 | 49.1 | 66.0 | 89.9 |
| Protein atoms | 59.2 | 38.6 | 29.6 | 59.8 | 68.1 | 48.8 | 66.3 | 90.2 |
| Protein main chain | 59.0 | 35.9 | 26.4 | 56.7 | 66.2 | 46.6 | 64.3 | 88.1 |
| Protein side chains | 59.4 | 41.2 | 32.7 | 62.9 | 70.1 | 50.9 | 68.2 | 92.3 |
| Inhibitor | - | - | - | - | 50.4 | 43.1 | 50.4 | 7.3 |
| Zn | 92.8 | 41.0 | 29.1 | 74.9 | 85.8 | 55.8 | 75.0 | 89.7 |
| Cl | 60.5 | 43.0 | 33.2 | 77.9 | 69.3 | 58.9 | 64.7 | 88.7 |
| Acetate | - | - | - | - | 52.5 | 51.3 | 49.5 | - |
| MES | - | - | - | - | 119.7 | 64.7 | 111.3 | 109.6 |
| PO<sub>4</sub> | 10 | 59.9 | 41.5 | 58.0 | - | - | - | - |
| Glycerol | - | 60.2 | 43.2 | - | - | - | - | - |
| Water | 32.7 | 45.1 | 42.6 | 50 | 59.6 | 51.4 | 59.1 | 71.7 |
| **Molprobit summary** |  |  |  |  |  |  |  |  |
| Ramachandran outliers (%) | 0.54 | 0.0 | 0.00 | 0.00 | 0.00 | 0.00 | 0.00 | 0.33 |
| Ramachandran favored (%) | 84.4 | 97.44 | 97.45 | 97.11 | 96.8 | 97.44 | 95.48 | 93.46 |
| Rotamer outliers (%) | 7.27 | 1.38 | 0.34 | 2.47 | 1.82 | 1.40 | 2.18 | 1.50 |
| C-beta outliers | 0 | 0 | 0 | 0 | 0 | 0 | 0 | 0 |
| Clashscore | 6.70 | 2.71 | 1.72 | 2.18 | 2.20 | 2.7 | 3.59 | 2.27 |
| MolProbity score | 2.69 | 1.28 | 1.04 | 1.45 | 1.39 | 1.28 | 1.72 | 1.57 |
| **PDB ID** | 6W9C | 6WZU | 6WRH | 6XG3 | 7JIR | 7JIT | 7JIV | 7JIW |

<sup>a</sup> Values in parentheses correspond to the highest resolution shell.
<sup>b</sup> $R_{\text{merge}} = \sum_h \sum_j |I_{hj} - \langle I_h \rangle| / \sum_h \sum_j I_{hj}$ , where $I_{hj}$ is the intensity of observation $j$ of reflection $h$ .
<sup>c</sup> As defined by Karplus and Diederichs <sup>37</sup>.
<sup>d</sup> $R = \sum_h |F_o| - |F_c| / \sum_h |F_o|$ for all reflections, where $F_o$ and $F_c$ are observed and calculated structure factors, respectively. $R_{\text{free}}$ is calculated analogously for the test reflections, randomly selected and excluded from the refinement.

## Supplemental Materials

### Materials and methods

#### Gene cloning, protein expression and purification of WT and C111S mutant of PLpro

The gene cloning, protein expression and purification were performed as reported previously^1^. Briefly, the Nsp3 DNA sequence corresponding to PLpro protease SARS-CoV-2 was optimized for *E. coli* expression using the OptimumGene codon optimization algorithm followed by manual editing and then cloned directly into pMCSG53 vector (Twist Bioscience). The plasmids were transformed into the *E. coli* BL21(DE3)-Gold strain (Stratagene). *E. coli* cells harboring plasmids for SARS CoV-2 PLpro WT and C111S mutant expression were cultured in LB medium supplemented with ampicillin (150 *µ*g/mL).

Bacterial cells were harvested by centrifugation at 7,000g and cell pellets were resuspended in a 12.5 ml lysis buffer (500 mM NaCl, 5% (v/v) glycerol, 50 mM HEPES pH 8.0, 20 mM imidazole pH8.0, 1mM TCEP, 1 *µ*M ZnCl₂) per liter culture and sonicated at 120W for 5 minutes (4 sec ON, 20 sec OFF). The cellular debris was removed by centrifugation at 30,000g for 90 minutes at 4 °C. The supernatant was mixed with 3 ml of Ni2+ Sepharose (GE Healthcare Life Sciences) which had been equilibrated with lysis buffer supplemented to 50 mM imidazole pH 8.0, and the suspension was applied on Flex-Column (420400-2510) connected to Vac-Man vacuum manifold (Promega). Unbound proteins were washed out via controlled suction with 160 ml of lysis buffer (with 50 mM imidazole pH 8.0). Bound proteins were eluted with 15 mL of lysis buffer supplemented to 500 mM imidazole pH 8.0, followed by Tobacco Etch Virus (TEV) protease treatment at 1:25 protease:protein ratio. The solutions were left at 4 °C overnight. The proteins were run separately on a Superdex 75 column equilibrated in lysis buffer. Fractions containing cut protein were collected and applied on Flex-Columns with 3 mL of Ni2+ Sepharose which had been equilibrated with lysis buffer. The flow through and a 7 mL lysis buffer rinse were collected. Lysis buffer was replaced using 30 kDa MWCO filters (Amicon-Millipore) via 10X concentration/dilution repeated 3 times to crystallization buffer (150 mM NaCl, 20 mM HEPES pH 7.5, 1 *µ*M ZnCl_2_, 4 mM TCEP). Purification was repeated for PLpro WT and C111S mutant proteins for co-crystallization with the inhibitor **3**, following the same protocol except that 10 mM β-mercaptoethanol was used instead of TCEP in all purification buffers, and 10 mM DTT was used instead of TCEP in the crystallization buffer. The final concentrations of WT PLpro was 25 mg/mL and C111S mutant was 30 mg/mL.

#### Fluorescence-based biochemical assays

Dose response assays were performed in 96-well plate format in triplicate at 25 °C. Wells containing varying concentrations of PLpro enzyme (0-1 *µ*M) in Tris-HCl pH 7.3, 1 mM EDTA were mixed with LKGG-AMC probe substrate (40 *µ*M) and measured continuously for fluorescence emission intensity (excitation λ: 364 nm; emission λ: 440 nm) on a Synergy Neo2 Hybrid. PLpro-WT and PLpro-C111S activities on LKGG-AMC were assayed as above with 1 *µ*M enzyme and 40 *µ*M LKGG-AMC substrate.

#### PLpro inhibition assay

Inhibition assays were performed in a 96-well plate format in triplicate at 25 °C. Reactions containing varying concentrations of inhibitor (0-500 *µ*M) and PLpro enzyme (0.3 *µ*M) in Tris-HCl pH 7.3, 1 mM EDTA were incubated for approximately five minutes. Reactions were then initiated with LKGG-AMC probe substrate (40 *µ*M), shaken linearly for 5 s, and then measured continuously for fluorescence emission intensity (excitation λ: 364 nm; emission λ: 440 nm) on a Synergy Neo2 Hybrid. Data were fit using nonlinear regression (dose-response inhibition, variable slope) analysis in GraphPad Prism 7.0.

#### PLpro crystallization

The sitting-drop vapor-diffusion method was used with the help of the Mosquito liquid dispenser (TTP LabTech) in 96-well CrystalQuick plates (Greiner Bio-One). Crystallizations were performed with the protein-to-matrix ratio of 1:1. MCSG1, MCSG2, MCSG3, and MCSG4 (Anatrace) screens were used for protein crystallization at 16 °C and 4 °C. The first thin-plate crystals grew in multiple conditions after 1-3 days of incubation at 4 °C. The best crystals of wild-type protein (C2 space group) were obtained from MCSG4 screen, reagent formulation #96 (0.2 M magnesium acetate, 10% PEG 8000). The crystals of PLpro-C111S mutant protein (bipyramidal crystals up to 0.2 mm, 1-3 days of incubation at 4 °C, P3_2_21 space group) were obtained from MCSG2 screen, reagent formulation #4 (0.1 M acetate buffer pH 4.5, 0.8 M NaH_2_PO_4_/1.2 M K_2_HPO_4_). The mutant crystals were used for seeding WT protein crystallization droplets to obtain crystals with significantly improved diffraction. For co-crystallization with inhibitors, proteins (15 mg/ml) were mixed with inhibitors at 10x protein concentration for a final inhibitor concentration of 4 mM, incubated on ice for 2.5 hours, and spun down to remove precipitation. Crystals (I4_1_22 space group) formed at 4 °C with a protein-to-matrix ratio of 2:1 in hanging drops in 0.1 M MES pH 6.0, 50 mM zinc acetate, 10% PEG 8000. Crystals selected for data collection were washed in the crystallization buffer supplemented with either 25% glycerol (apo-protein crystals) or 25% ethylene glycol (protein-inhibitor crystals) and flash-cooled in liquid nitrogen.

#### Data collection, structure determination and refinement

Single-wavelength x-ray diffraction data were collected at 100 K temperature at the 19-ID beamline of the Structural Biology Center at the Advanced Photon Source at Argonne National Laboratory using the program SBCcollect. The diffraction images were recorded from all crystal forms on the PILATUS3 X 6M detector using 0.3° rotation and 0.3 sec exposure (with the exception of the first WT PLpro crystal where data were collected using 0.5° rotation and 0.5 sec exposure). The intensities were integrated and scaled with the HKL3000 suite ^2^. Intensities were converted to structure factor amplitudes in the truncate program ^3,4^ from the CCP4 package ^5^. The structures were determined by molecular replacement using HKL3000 suite incorporating the following programs: MOLREP ^6^, SOLVE/RESOLVE ^7^ and ARP/wARP ^8^. The initial solutions were refined, both rigid-body refinement and regular restrained refinement by REFMAC program ^9^ as a part of HKL3000. The coordinates of SARS coronavirus PLpro (PDB id: 5Y3Q) were used as the starting model for the first wild-type protein structure solution. Several rounds of manual adjustments of structure models using COOT ^10^ and refinements with REFMAC program ^9^ from CCP4 suite ^5^ were done. The models including the ligands were manually adjusted using COOT and then iteratively refined using COOT and REFMAC. The stereochemistry of the structure was validated with PHENIX suite ^11^ incorporating MOLPROBITY ^12^ tools. Throughout the refinement, the same 5% of reflections were kept out from the refinement. The stereochemistry of the structure was checked with PROCHECK ^13^ and the Ramachandran plot and validated with the PDB validation server. A summary of data collection and refinement statistics is given in Supplemental Table 1.

### Chemical synthesis methods

^1^H NMR and ^13^C NMR spectra were collected at 25 °C on 400 MHz Bruker DRX400 at the Department of Chemistry NMR Facility at the University of Chicago. ^1^H-NMR chemical shifts are reported in parts per million (ppm) relative to the peak of residual proton signals from (CDCl_3_ 7.26 ppm or DMSO-*d*_6_ 2.50 ppm). Multiplicities are given as: s (singlet), d (doublet), t (triplet), q (quartet), dd (doublet of doublets), p (pentet), m (multiplet), and br (broad) ^14^. ^13^C-NMR chemical shifts are reported in parts per million (ppm) relative to the peak of residual proton signals from (CDCl_3_ 77.16 ppm). Analysis of NMR was done in MestReNova (version 14.1.2-25024). High resolution mass was obtained from Agilent 6224 TOF High Resolution Accurate Mass Spectrometer (HRA-MS) using combination of APCI and ESI at the Department of Chemistry Mass Spectrometry Facility at the University of Chicago. Low resolution mass spectral analyses and liquid chromatography analysis were carried out on an Advion Expression-L mass spectrometer (Ithaca, NY) coupled with an Agilent 1220 Infinity LC System (Santa Clara, CA).

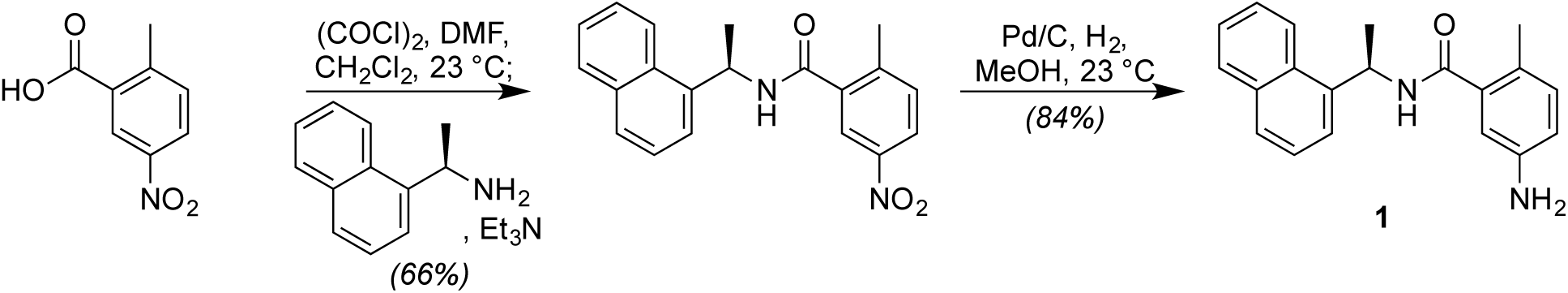

### Synthesis of Compound 1 (GRL0617)

To a suspension of the 5-nitro-*o*-toluic acid (0.362 g, 2.00 mmol, 1.0 equiv) in CH_2_Cl_2_ (10.0 mL) at 0 °C was added (COCl)_2_ (0.206 mL, 2.40 mmol, 1.2 equiv) dropwise before catalytic DMF (8 drops from a 1.00 mL syringe) was added dropwise. This mixture was then stirred at 0 °C for 30 min before being concentrated directly. The resultant residue was then redissolved in CH_2_Cl_2_ (20.0 mL) at 23 °C before the sequential addition of (*R*)-(+)-1-(1-naphthyl)ethylamine (0.417 mL, 2.60 mmol, 1.3 equiv) and Et_3_N (0.558 mL, 4.00 mmol, 2.0 equiv). The reaction mixture was then stirred at 23 °C for 30 min. Upon completion, the reaction contents were quenched by the addition of 1 M HCl (30 mL), diluted with CH_2_Cl_2_ (10 mL), and poured into a separatory funnel. The two phases were separated, and the organic layer was washed with 1 M NaOH (3 × 30 mL). The organic extract was then dried (Na_2_SO_4_), filtered, and concentrated. Purification of the resultant residue by flash column chromatography (silica gel, hexanes:EtOAc, 2:1) afforded the desired amide intermediate (0.440 g, 66% yield) as a white solid. Pd/C (0.200 g, 10% by weight, 0.19 mmol, 0.15 equiv) was then carefully added to a suspension of the newly formed amide intermediate (0.430 g, 1.29 mmol, 1.0 equiv) in MeOH (30.0 mL) at 23 °C. The resultant suspension was then purged by direct bubbling with a balloon of H_2_ gas for 2 h at 23 °C. Upon completion, the reaction contents were filtered through a short pad of Celite, washed with EtOAc (200 mL), and dried (Na_2_SO_4_) to directly provide inhibitor **1** (0.328 g, 84% yield) as a white solid. **1 (GRL0617)**: R*_f_* = 0.20 (silica gel, hexanes:EtOAc, 1:1); IR (film) ν_max_ 3339, 3049, 2976, 2927, 1639, 1511, 1339, 1244, 1121, 817, 800, 755 cm^−1^; ^1^H NMR (400 MHz, CDCl_3_) δ 8.23 (d, *J* = 8.7 Hz, 1 H), 7.88 (d, *J* = 8.9 Hz, 1 H), 7.81 (d, *J* = 8.1 Hz, 1 H), 7.60–7.49 (m, 3 H), 7.46 (dd, *J* = 8.2, 7.2 Hz, 1 H), 6.93 (d, *J* = 7.9 Hz, 1 H), 6.60–6.54 (m, 2 H), 6.16–6.07 (m, 1 H), 5.97 (d, *J* = 8.5 Hz, 1 H), 3.51 (s, 2 H), 2.29 (s, 3 H), 1.77 (d, *J* = 6.7 Hz, 3 H); ^13^C NMR (101 MHz, CDCl_3_) δ 169.2, 144.2, 138.2, 137.1, 134.1, 131.9, 131.3, 128.9, 128.6, 126.7, 126.1, 125.5, 125.3, 123.7, 122.7, 116.8, 113.5, 44.9, 20.8, 18.9; [α]_D_^22^ = –75.8° (*c* = 1.0, CHCl_3_) [lit. [α]_D_^20^ = –76.8° (*c* = 1.0, CHCl_3_) from *J. Med. Chem.* **2009**, *52*, 5228].

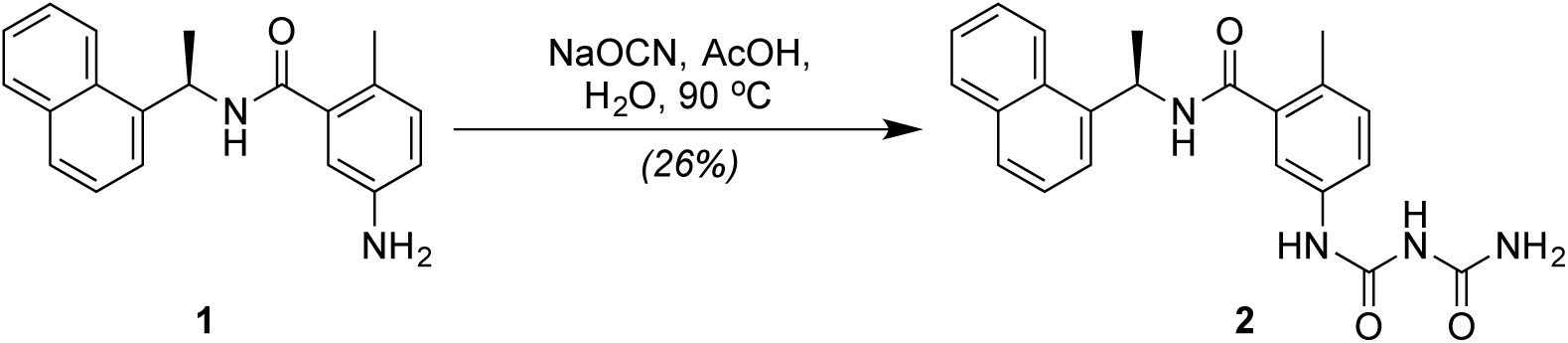

### Synthesis of Compound 2

To a solution of amine **1** (30.0 mg, 0.10 mmol, 1.0 equiv) in CH_3_CN/H_2_O (1:1) (1.2 mL) was added NaOCN (51.3 mg, 0.80 mmol, 8.0 equiv) and the mixture was heated at 90 °C for 15 min. Then, AcOH (23.2 μL, 24.2 mg, 0.40 mmol, 4.0 equiv) was added dropwise, and the stirring was continued for additional 1 h at 90 °C. Then, the second portion of AcOH (23.2 μL, 24.2 mg, 0.40 mmol, 4.0 equiv) was introduced, and the resulting solution was stirred for 3 h at 90 °C. Upon completion, the mixture was cooled to 23 °C, the precipitate filtered, washed with H_2_O (5 × 2 mL) and dried (using air) to afford compound **2** (10.6 mg, 26% yield) as a white solid. **2**: R*_f_* = 0.20 (silica gel, EtOAc). IR (film) ν_max_ 3284, 2975, 2928, 1704, 1640, 1548, 1496, 1408, 1228, 1201, 801, 779 cm^−1^; ^1^H NMR (400 MHz, DMSO-*d*_6_) δ 9.97 (s, 1H), 8.90 (d, *J* = 8.0 Hz, 1 H), 8.86–8.84 (m, 1 H), 8.24 (d, *J* = 8.4 Hz, 1 H), 7.98–7.92 (m, 1 H), 7.87–7.80 (m, 1 H), 7.63–7.49 (m, 4 H), 7.43–7.35 (m, 2 H), 7.15 (d, *J* = 8.2 Hz, 1 H), 7.05–6.71 (m, 2 H), 5.91 (p, *J* = 7.1 Hz, 1 H), 2.22 (s, 3 H), 1.57 (d, *J* = 6.9 Hz, 3 H); ^13^C NMR (101 MHz, DMSO-*d*_6_): δ 167.8, 155.5, 152.0, 140.2, 137.6, 135.6, 133.4, 130.8, 130.4, 129.5, 128.7, 127.3, 126.2, 125.6, 125.5, 123.2, 122.5, 119.8, 117.7, 44.3, 21.5, 18.6; HRMS (ESI) calculated for C_22_H_23_N_4_O_3_^+^ [M + H^+^] 391.1765, found 391.1761; [α]_D_^22^ = –102.3° (*c* = 0.2, acetone).

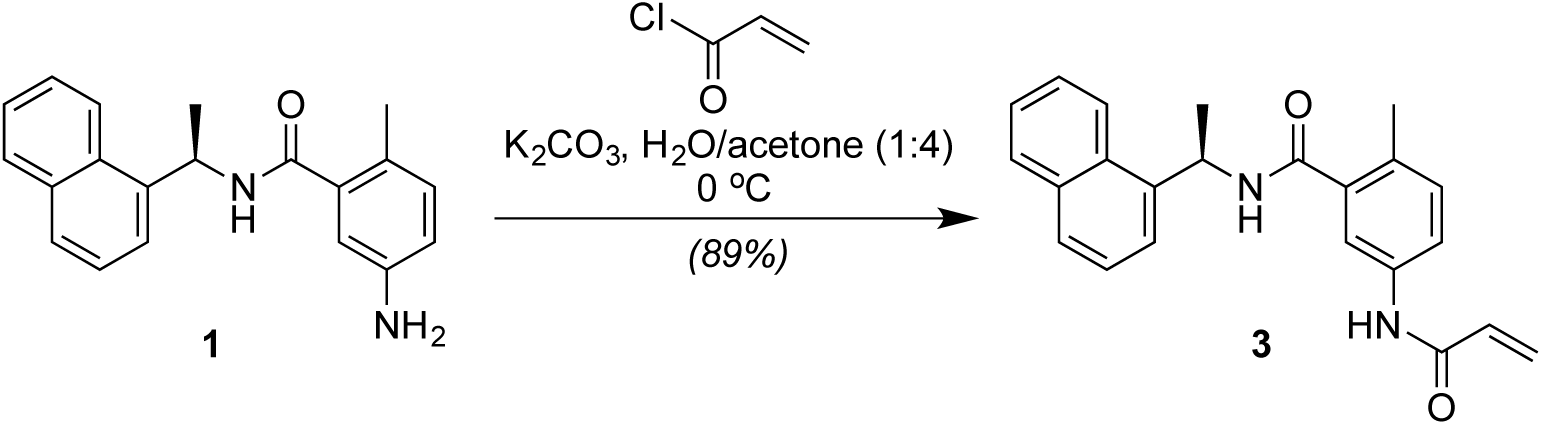

### Synthesis of Compound 3

To a solution of K_2_CO_3_ (18.2 mg, 0.132 mmol, 2.0 equiv) in H_2_O (0.12 mL) and acetone (0.48 mL) at 0 °C was added acryloyl chloride (0.011 mL, 0.132 mmol, 2.0 equiv). Amine **1** (20.0 mg, 0.066 mmol, 1.0 equiv) was then added dropwise at 0 °C as a solution in acetone (0.2 mL) and the reaction mixture was allowed to stir at 0 °C for 10 min. Upon completion, the reaction contents were quenched by the addition of saturated aqueous NH_4_Cl (10 mL), diluted with CH_2_Cl_2_ (10 mL), and poured into a separatory funnel. The two phases were separated, and the aqueous layer was extracted with CH_2_Cl_2_ (3 × 10 mL). The combined organic extracts were then dried (Na_2_SO_4_), filtered, and concentrated. Purification of the resultant residue by flash column chromatography (silica gel, hexanes:EtOAc, 1:1) afforded the desired acrylamide **3** (21.0 mg, 89% yield) as a white solid. **3**: R*_f_* = 0.27 (silica gel, hexanes:EtOAc, 1:1); IR (film) ν_max_ 3276, 3052, 2977, 1707, 1611, 1596, 1541, 1496, 1411, 1244, 1202, 982, 799, 778 cm^−1^; ^1^H NMR (400 MHz, CDCl_3_) δ 8.19 (d, *J* = 8.3 Hz, 1 H), 8.06 (s, 1 H), 7.85 (dd, *J* = 7.9, 1.7 Hz, 1 H), 7.77 (d, *J* = 8.2 Hz, 1 H), 7.56–7.45 (m, 3 H), 7.44–7.37 (m, 2 H), 7.32 (s, 1 H), 6.99 (d, *J* = 8.3 Hz, 1 H), 6.43 (d, *J* = 8.3 Hz, 1 H), 6.34 (dd, *J* = 16.9, 1.5 Hz, 1 H), 6.19 (dd, *J* = 16.9, 10.1 Hz, 1 H), 6.12–6.02 (m, 1 H), 5.66 (dd, *J* = 10.1, 1.5 Hz, 1 H), 2.29 (s, 3 H), 1.73 (d, *J* = 6.8 Hz, 3 H); ^13^C NMR (110 MHz, CDCl_3_) δ 169.0, 163.9, 138.2, 136.5, 135.5, 134.1, 132.1, 131.6, 131.2, 131.1, 129.0, 128.6, 127.9, 126.7, 126.0, 125.4, 123.5, 122.8, 122.0, 118.8, 45.2, 21.0, 19.3; HRMS (ESI) calculated for C_23_H_23_N_2_O_2_^+^ [M + H^+^] 359.1754, found 359.1754; [α]_D_^22^ = –110.7° (*c* = 1.0, acetone). [Note: a slight concentration dependence was observed for NMR spectra of this compound].

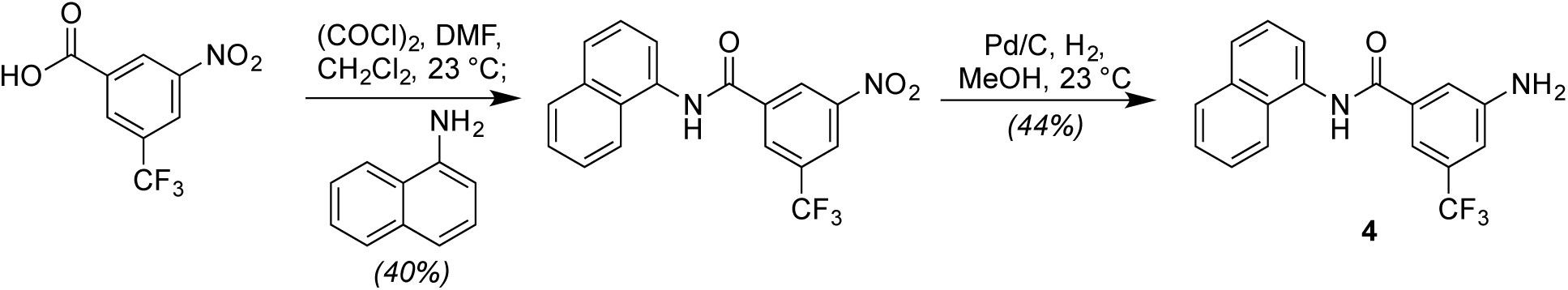

### Synthesis of Compound 4

3-nitro-5-(trifluoromethyl)benzoic acid (0.752 g, 3.20 mmol, 1.0 equiv) was suspended in CH_2_Cl_2_ (14 mL) under an argon atmosphere at 23 °C. Then, DMF (2 drops) was added, followed by slow addition of oxalyl chloride (0.300 mL, 0.444 g, 3.50 mmol, 1.1 equiv). The resulting solution was stirred at 23 °C for 1 h before complete dissolution of the starting material was observed. The mixture was then concentrated to dryness, back-filled with argon, redissolved in CH_2_Cl_2_ (10 mL) and cooled to 0 °C. Then, naphthylamine (0.500 g, 3.49 mmol, 1.1 equiv) was added in a single portion and the solution was warmed to 23 °C and stirred vigorously at this temperature for 2 h. The resulting precipitate was filtered and washed with cold CH_2_Cl_2_ (2 × 10 mL). The CH_2_Cl_2_-containing filtrate was discarded, and the filter cake was then thoroughly washed with warm (50 °C) EtOAc (4 × 20 mL). The filtrate was concentrated to afford the desired amide (0.460 g, 40% yield) as a white solid. R*_f_* = 0.36 (silica gel, hexanes:EtOAc = 5:1); ^1^H NMR (400 MHz, DMSO-*d*_6_) δ 11.00 (s, 1 H), 9.19–9.12 (m, 1 H), 8.88 (s, 1 H), 8.78–8.73 (m, 1 H), 8.09–7.99 (m, 2 H), 7.96–7.87 (m, 1 H), 7.69–7.53 (m, 4 H). Next, to a suspension of the newly formed amide (0.050 g, 0.14 mmol, 1.0 equiv) in MeOH/EtOAc (2 mL, 1:1) at 23 °C was flushed several times with nitrogen and then charged with Pd/C (10 wt %, 20 mg). After flushing the resulting solution several times with H_2_, the reactions contents were left to stir at 23 °C under a H_2_ atmosphere for 12 h. Upon completion, the resulting solution was filtered through Celite^®^ (washing with MeOH) and concentrated. The resulting crude product was suspended in CH_2_Cl_2_ (2 mL) followed by addition of methanol (with stirring) until a clear solution was obtained. The resulting mixture was placed in the freezer (–20 °C) overnight. The precipitate was then collected by filtration, rinsed with CH_2_Cl_2_ and dried on high vacuum to afford compound **4** (20.0 mg, 44% yield) as a white solid. **4**: R*_f_* = 0.30 (silica gel, hexanes:EtOAc, 1:1); IR (film) ν_max_ 3355, 3229, 3053, 1627, 1605, 1526, 1504, 1371, 1264, 1168, 1122, 998, 867, 691 cm^−1^; ^1^H NMR (400 MHz, DMSO-*d*_6_) δ 10.47 (s, 1 H), 8.03–7.92 (m, 2 H), 7.91–7.84 (m, 1 H), 7.61–7.45 (m, 6 H), 7.11–7.00 (m, 1 H), 5.93–5.82 (br s, 2 H); ^13^C NMR (101 MHz, DMSO-*d*_6_) δ 165.7, 149.8, 136.3, 133.8, 129.9 (q, *J* = 31.3 Hz), 129.2, 128.1, 126.4, 126.1, 126.0, 125.7, 125.6, 124.0, 123.3, 123.0, 116.7, 112.1 (q, *J* = 4.0 Hz), 110.6 (q, *J* = 3.9 Hz); ^19^F NMR (470 MHz, DMSO-*d*_6_) δ –61.34; HRMS (ESI) calculated for C_18_H_14_F_3_N_2_O^+^ [M + H^+^] 331.1053, found 331.1052.

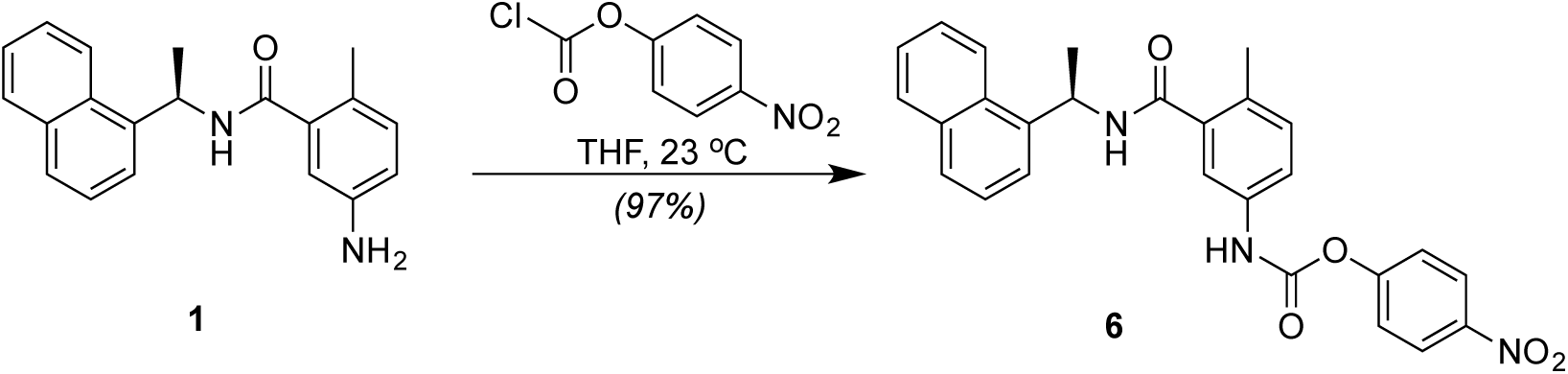

### Synthesis of Compound 6

To a solution of amine **1** (21.9 mg, 0.072 mmol, 1.0 equiv) in THF (0.70 mL) at 23 °C was added 4-nitrophenyl chloroformate (21.8 mg, 0.108 mmol, 1.5 equiv) and the reaction mixture was allowed to stir at 23 °C for 1 h. Upon completion, the solution was concentrated directly. Purification of the resultant residue by flash column chromatography (silica gel, hexanes:EtOAc, 2:1) afforded compound **6** (32.4 mg, 97% yield) as a white solid. **6**: R*_f_* = 0.27 (silica gel, hexanes:EtOAc, 1:1); IR (film) ν_max_ 3254, 3051, 2051, 1735, 1638, 1603, 1523, 1489, 1345, 1201, 1011, 858, 778 cm^−1^; ^1^H NMR (500 MHz, CDCl_3_) δ 8.19–8.16 (m, 1 H), 8.15–8.12 (m, 2 H), 7.88–7.83 (m, 1 H), 7.80–7.77 (m, 1 H), 7.77–7.74 (m, 1 H), 7.54–7.46 (m, 3 H), 7.45–7.38 (m, 2 H), 7.35–7.29 (m, 1 H), 7.20–7.14 (m, 2 H), 7.09 (d, *J* = 8.3 Hz, 1 H), 6.26–6.21 (m, 1 H), 6.10 (p, *J* = 6.8 Hz, 1 H), 2.35 (s, 3 H), 1.74 (d, *J* = 6.8 Hz, 3 H). ^13^C NMR (126 MHz, CDCl_3_) δ 168.7, 155.3, 150.5, 148.4, 145.0, 137.8, 137.0, 134.9, 134.1, 131.8, 131.2, 129.0, 128.7, 126.7, 126.1, 125.3, 125.2, 123.4, 122.7, 122.2, 120.7, 117.7, 45.2, 20.8, 19.2; HRMS (ESI) calculated for C_27_H_24_N_3_O_5_^+^ [M + H^+^] 470.1710, found 470.1703; [α]_D_^20^ = –78.4° (*c* = 1.0, CHCl_3_).

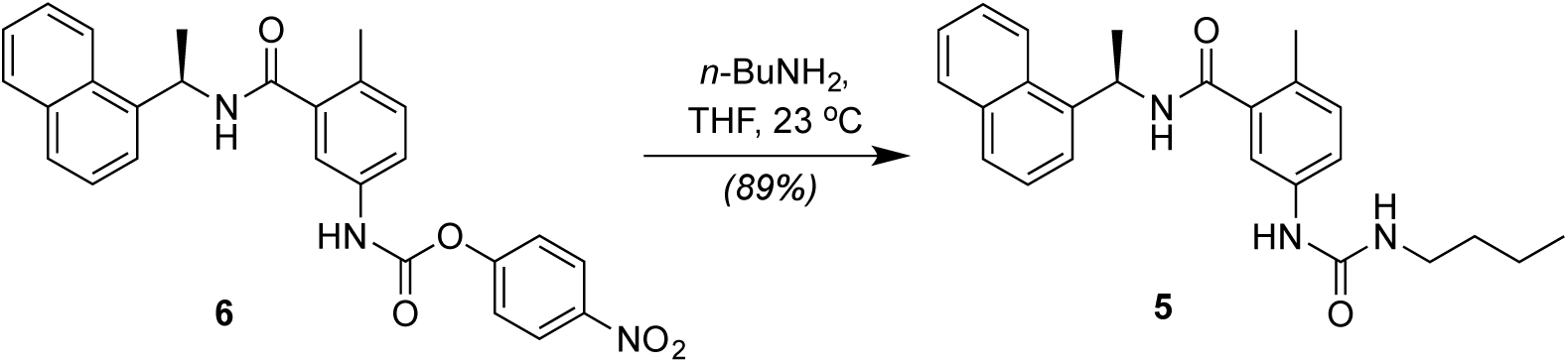

### Synthesis of Compound 5

To a solution of **6** (8.5 mg, 0.018 mmol, 1.0 equiv) in THF (0.20 mL) at 23 °C was added *n*-butylamine (0.01 mL, 0.036 mmol, 2.0 equiv) and the reaction mixture was allowed to stir for 5 min. Upon completion, the reaction contents were quenched by the addition of H_2_O (5 mL), diluted with CH_2_Cl_2_ (5 mL), and poured into a separatory funnel. The two phases were separated, and the aqueous layer was extracted with CH_2_Cl_2_ (3 × 30 mL). The organic extract was then dried (Na_2_SO_4_), filtered, and concentrated. Purification of the resultant residue by flash column chromatography (silica gel, CH_2_Cl_2_:MeOH, 40:1) afforded compound **5** (6.3 mg, 89% yield) as a white solid. **5**: R*_f_* = 0.41 (silica gel, CH_2_Cl_2_:MeOH, 40:1); IR (film) ν_max_ 3276, 2978, 2946, 2738, 2603, 1633, 1548, 1445, 1236, 1172, 1037, 806 cm^−1^; ^1^H NMR (400 MHz, CDCl_3_) δ 8.21 (d, *J* = 8.4 Hz, 1 H), 7.88 (d, *J* = 7.7 Hz, 1 H), 7.80 (d, *J* = 8.2 Hz, 1 H), 7.58–7.49 (m, 3 H), 7.45 (dd, *J* = 8.2, 7.2 Hz, 1 H), 7.21–7.15 (m, 2 H), 7.07 (d, *J* = 8.2 Hz, 1 H), 6.25 (s, 1 H), 6.11 (d, *J* = 5.6 Hz, 2 H), 4.66 (s, 1 H), 3.17 (q, *J* = 6.7 Hz, 2 H), 2.34 (s, 3 H), 1.77 (d, *J* = 6.3 Hz, 3 H), 1.47–1.40 (m, 2 H), 1.31 (dd, *J* = 15.0, 7.5 Hz, 2 H), 0.90 (t, *J* = 7.3 Hz, 3 H); ^13^C NMR (126 MHz, DMSO-*d*_6_) δ 168.2, 155.2, 140.3, 138.1, 137.4, 133.4, 130.5, 128.7, 127.6, 127.2, 127.0, 126.2, 125.6, 125.5, 123.2, 122.5, 118.3, 116.2, 44.2, 38.7, 31.9, 21.5, 19.5, 18.5, 13.7; HRMS (ESI) calculated for C_25_H_30_N_3_O_2_^+^ [M + H^+^] 404.2333, found 404.2337; [α]_D_^20^ = –32.5° (*c* = 0.4, acetone).

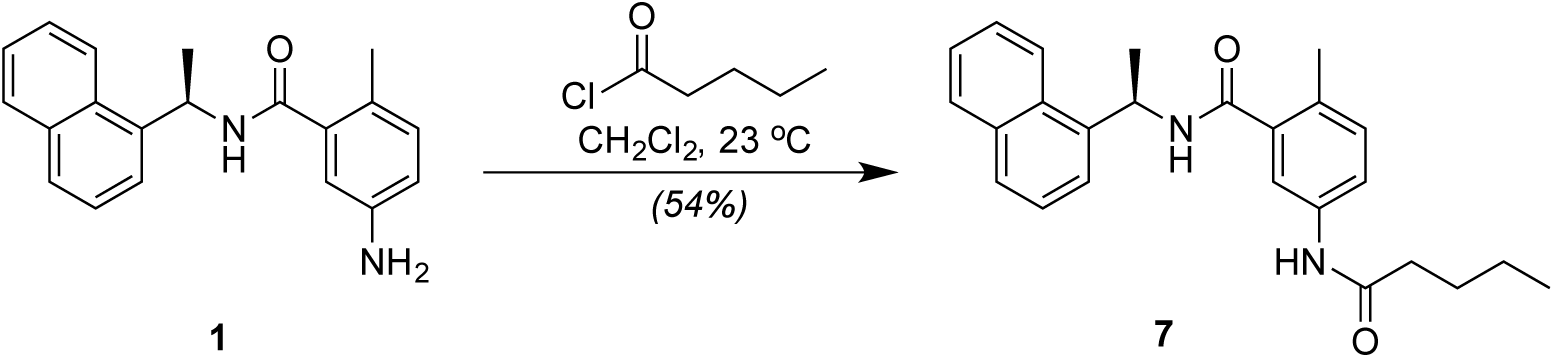

### Synthesis of Compound 7

To a solution of amine **1** (24.4 mg, 0.080 mmol, 1.0 equiv) in CH_2_Cl_2_ (0.800 mL) at 23 °C was added valeroyl chloride (0.010 mL, 0.083 mmol, 1.0 equiv) dropwise. Upon completion, the reaction contents were quenched by the addition of H_2_O (1 mL), diluted with CH_2_Cl_2_ (1 mL), and poured into a separatory funnel. The two phases were separated, and the aqueous layer was extracted with CH_2_Cl_2_ (3 × 1 mL). The combined organic extracts were then dried (Na_2_SO_4_), filtered, and concentrated. Purification of the resultant residue by flash column chromatography (silica gel, hexanes:EtOAc, 1:1) afforded compound **7** (16.8 mg, 54% yield) as a colorless oil. **7**: R*_f_* = 0.50 (silica gel, hexanes:EtOAc, 1:1); IR (film) ν_max_ 3274, 3051, 2959, 2929, 2871, 1638, 1594, 1541, 1452, 1338, 1188, 1091, 823, 799, 777 cm^−1^; ^1^H NMR (400 MHz, CDCl_3_) δ 8.22 (d, *J* = 8.4 Hz, 1 H), 7.87 (d, *J* = 8.0 Hz, 1 H), 7.80 (d, *J* = 8.2 Hz, 1 H), 7.59–7.41 (m, 5 H), 7.37 (d, *J* = 8.2 Hz, 1 H), 7.29 (s, 1 H), 7.08 (d, *J* = 8.2 Hz, 1 H), 6.18 (d, *J* = 8.4 Hz, 1 H), 6.10 (app p, *J* = 6.8 Hz, 1 H), 2.36 (s, 3 H), 2.28 (t, *J* = 7.6 Hz, 2 H), 1.77 (d, *J* = 6.8 Hz, 3 H), 1.65 (p, *J* = 7.6 Hz, 2 H), 1.36 (hex, *J* = 7.4 Hz, 2 H), 0.92 (t, *J* = 7.4 Hz, 3 H); ^13^C NMR (101 MHz, CDCl_3_) δ 171.7, 168.7, 138.1, 136.8, 135.8, 134.1, 131.9, 131.6, 131.3, 129.0, 128.6, 126.7, 126.1, 125.4, 123.6, 122.8, 121.6, 118.4, 45.1, 37.5, 27.7, 22.5, 20.9, 19.4, 13.9.; HRMS (ESI) calculated for C_25_H_29_N_2_O_2_^+^ [M + H^+^] 389.2224, found 389.2220; [α]_D_^20^ = –81.7° (*c* = 0.3, CHCl_3_).

### Synthesis of S1

A solution of Boc glycine (507.8 mg, 2.89 mmol, 1.0 equiv), oxyma (0.540 g, 3.79 mmol, 1.2 equiv), DIC (0.538 *µ*L, 3.47 mmol, 1.2 eq) and fluorophore amine (0.389 g, 2.22 mmol, 0.75 eq) in dry DMF (15 mL) was first stirred at 35 °C for 1 h and then overnight at 45 °C. The reaction mixture was concentrated upon reduced pressure and purification by column chromatography (silica gel, 2-10% MeOH:CH_2_Cl_2_) yielded **S1** in 78% purity as analyzed by LCMS. For further purification, the impure mixture was dissolved in minimal CH_2_Cl_2_, washed with water and brine. The organic layer was cooled to 0 °C and the white precipitate was filtered to yield pure **S1** (0.562 g, 76% yield). **S1**: ^1^H NMR (400 MHz, DMSO-*d*_6_) δ 10.39 (s, 1 H), 7.78–7.69 (m, 2 H), 7.48 (dd, *J* = 8.7, 2.0 Hz, 1 H), 7.12 (t, *J* = 6.1 Hz, 1 H), 6.26 (d, *J* = 1.3 Hz, 1 H), 3.77 (d, *J* = 6.1 Hz, 2 H), 2.40 (d, *J* = 1.2 Hz, 3 H), 1.40 (s, 9 H); LRMS (ESI) calculated for C_17_H_21_N_2_O_5_ [M + H^+^] 333.14, found 333.4. ^15^

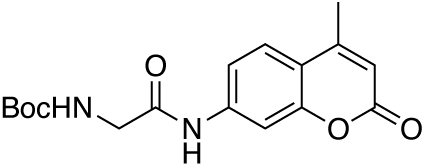

### Synthesis of S2

EDC•HCl (5.68 g, 29.5 mmol, 1.3 equiv) was added to a solution of Fmoc-Lys(Boc)-OH (10.02 g, 22.7 mmol, 1.0 equiv), HOBt (4.34 g, 23.8 mmol, 1.05 equiv), HCl•H_2_N-Gly-OtBu (3.80 g, 22.7 mmol, 1.0 equiv) and *i-*Pr_2_NEt (5.93 mL, 34.05 mmol, 1.5 equiv) in DMF (50 mL) at 23 °C. The resulting reaction mixture was stirred for 16 h at 23 °C and then concentrated under reduced pressure. Purification by column chromatography (silica gel, 30-70% EtOAc:hexane) yielded **S2** (11.3 g, 86% yield). **S2**: ^1^H NMR (400 MHz, CDCl_3_) δ 7.76 (d, *J* = 7.5 Hz, 2 H), 7.59 (d, *J* = 7.7 Hz, 2 H), 7.39 (t, *J* = 7.5 Hz, 2 H), 7.31 (td, *J* = 7.4, 1.2 Hz, 2 H), 6.53 (s, 1 H), 5.52 (s, 1 H), 4.66 (s, 1 H), 4.40 (d, *J* = 8.2 Hz, 2 H), 4.21 (t, *J* = 6.9 Hz, 2 H), 3.92 (s, 2 H), 3.11 (d, *J* = 5.3 Hz, 2 H), 1.97–1.80 (m, 1 H), 1.60–1.75 (m, 2 H), 1.30–1.55 (m, 21 H); LRMS (ESI) calculated for C_32_H_44_N_3_O_7_ [M + H^+^] 582.31, found 582.6.^16^

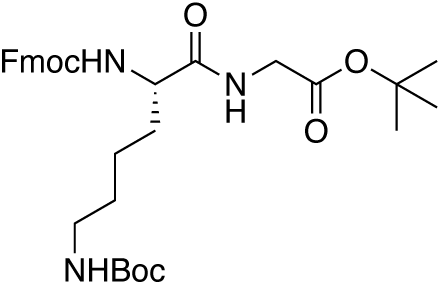

### Synthesis of S3

A solution of **S2** (1.392 g, 2.39 mmol, 1.0 equiv) in 10% Piperidine:CH_2_Cl_2_ (20 mL) was stirred for 20 min at 23 °C and followed with evaporation by rotary evaporation. Dilution with DMF (20 mL) and evaporation by rotary evaporation was repeated two more times to remove residual piperidine. The resultant crude was resuspended in DMF (30 mL), Fmoc leucine (0.928g, 2.62 mmol, 1.1 equiv), HOBt (80%, 0.445 g, 2.63 mmol, 1.2 equiv) and EDC•HCl (0.596 g, 3.10 mmol, 1.3 equiv) were added then added at 23 °C. The reaction mixture was stirred for 30 min at 23 °C and concentrated under reduced pressure. Purification by column chromatography (silica gel, 10-50% EtOAc:CH_2_Cl_2_) yielded **S3** (1.659 g, 85% yield). **S3**: ^1^H NMR (400 MHz, CDCl_3_) δ 7.74 (d, *J* = 7.6 Hz, 2 H), 7.58 (d, *J* = 7.4 Hz, 2 H), 7.38 (t, *J* = 7.5 Hz, 2 H), 7.29 (dd, *J* = 7.5, 3.4 Hz, 2 H), 6.93 (s, 1 H), 6.84 (d, *J* = 7.1 Hz, 1 H), 5.68 (s, 1 H), 4.81 (s, 1 H), 4.62–4.48 (m, 1 H), 4.42 (dd, *J* = 10.6, 7.2 Hz, 1 H), 4.34 (t, *J* = 8.8 Hz, 1 H), 4.25 (d, *J* = 5.1 Hz, 1 H), 4.19 (t, *J* = 7.1 Hz, 1 H), 3.99–3.80 (m, 2 H), 3.03 (d, *J* = 6.6 Hz, 2 H), 2.01 (s, 1 H), 1.87 (h, *J* = 7.6 Hz, 1 H), 1.74–1.51 (m, 4 H), 1.43 (s, 9 H), 1.40 (s, 9 H), 1.38–1.22 (m, 3 H), 0.92 (d, *J* = 6.5 Hz, 6 H); ^13^C NMR (101 MHz, CDCl_3_) δ 172.72, 171.60, 168.84, 156.56, 143.96, 143.82, 141.39, 127.84, 127.19, 125.24, 120.10, 82.41, 77.48, 76.84, 67.20, 53.76, 52.89, 47.24, 42.11, 41.52, 32.06, 29.50, 28.56, 28.14, 24.81, 23.13, 22.62, 21.96; HRMS (ESI) calculated for C_38_H_54_N_4_O_8_ [M^+^] 694.3942, found 694.3945.

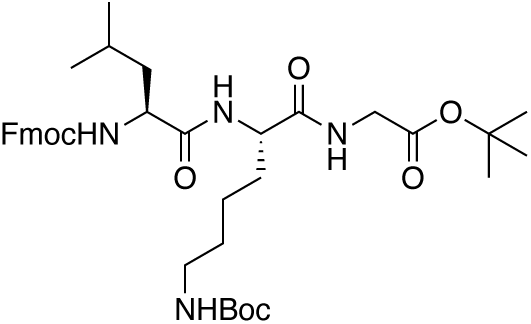

### Synthesis of S4

A solution of **S3** (1.23 g, 1.77 mmol, 1.0 equiv) and Et_3_SiH (1.4 mL, 8.76 mmol, 5 equiv) in 25% TFA:CH_2_Cl_2_ (28 mL) was stirred at 23 °C for 18 h, followed by dilution with CH_2_Cl_2_ (20 mL) and solvent removal by rotary evaporation. Dilution with CH_2_Cl_2_ (30 mL) and solvent evaporation by rotary evaporation was repeated two more times to remove residual TFA. The resultant crude material was resuspended in MeOH (15 mL) followed by addition of Boc_2_O (0.511 g, 2.22 mmol, 1.25 equiv) and *i*-Pr_2_NEt (2.29 mL, 13.1 mmol, 7.5 equiv). The reaction mixture was stirred for 1 h at 23 °C, diluted with EtOAc (100 mL), and washed with aqueous HCl (pH ∼2). The aqueous layer was further extracted by EtOAc (2 × 30 mL), the combined organic layers were dried over Na_2_SO_4_, and then evaporated to give a crude product. Purification by column chromatography (silica gel, 3-15% MeOH:CH_2_Cl_2_) yielded **S4** (0.781 g, 87% purity by LCMS). **S4**: HRMS (ESI) calculated for C_34_H_46_N_4_O_8_ [M^+^] 638.3316, found 638.3312

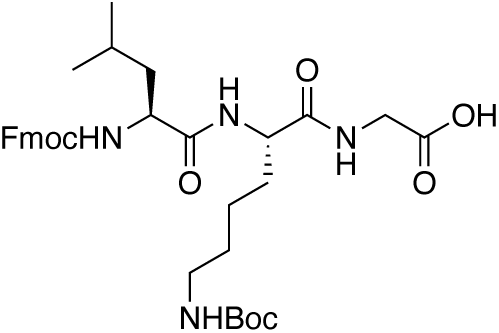

### Synthesis of S5

A solution of **S1** (47.8 mg, 0.143 mmol) in 20% TFA:CH_2_Cl_2_ (2 mL) was stirred at 23 °C for 30 min, followed by dilution with CH_2_Cl_2_ (10 mL) and solvent removal by rotary evaporation. Dilution with CH_2_Cl_2_ (15 mL) and solvent removal by rotary evaporation was repeated two more times to remove residual TFA. The resultant crude was resuspended in DMF (5 mL) and deprotected amine intermediate (2.7 mL, 77.4 *µ*mol, 1.0 equiv) was added to a flask containing **S4** (49.7 mg, 77.8 *µ*mol, 1.0 equiv) and *i*-Pr_2_NEt (81.0 *µ*L, 0.46 mmol, 6.0 equiv) in DMF (1 mL) at 23 °C. After the addition of EDC•HCl (31.4 mg, 0.15 mmol, 2.0 equiv) the reaction mixture was stirred for 16 h at 23 °C, diluted by EtOAc (20 mL), and then washed with aqueous HCl (pH ∼2-3). The aqueous layer was further extracted by EtOAc (2 × 20 mL), the combined organic layers were dried over Na_2_SO_4_, and then evaporated to give a crude product. Purification by column chromatography (silica gel, 1-6% MeOH in 1:1 EtOAc:CH_2_Cl_2_) yielded **S5** (22.7 mg, 34% yield). **S5**: ^1^H NMR (400 MHz, DMSO-*d*_6_) δ 10.15 (s, 1 H), 8.50–8.38 (m, 1 H), 8.26 (t, *J* = 5.9 Hz, 1 H), 8.07 (d, *J* = 6.8 Hz, 1 H), 7.88 (d, *J* = 7.6 Hz, 2 H), 7.79 (d, *J* = 2.0 Hz, 1 H), 7.75–7.66 (m, 3 H), 7.56–7.47 (m, 2 H), 7.40 (t, *J* = 6.9 Hz, 2 H), 7.30 (tdd, *J* = 7.4, 2.2, 1.2 Hz, 2 H), 6.71 (d, *J* = 6.7 Hz, 1 H), 6.27 (d, *J* = 1.3 Hz, 1 H), 4.36–4.14 (m, 4 H), 4.13–4.02 (m, 2 H), 3.95 (d, *J* = 5.9 Hz, 2 H), 3.84–3.71 (m, 2 H), 3.17 (d, *J* = 5.2 Hz, 1 H), 2.87 (d, *J* = 5.9 Hz, 2 H), 2.39 (d, *J* = 1.3 Hz, 3 H), 1.55 (td, *J* = 41.8, 8.0 Hz, 4 H), 1.34 (s, 12 H), 0.79 (dd, *J* = 10.8, 6.6 Hz, 6 H); HRMS (ESI) calculated for C_46_H_56_N_6_O_10_Na [M + Na^+^] 875.3956, found 875.3944.

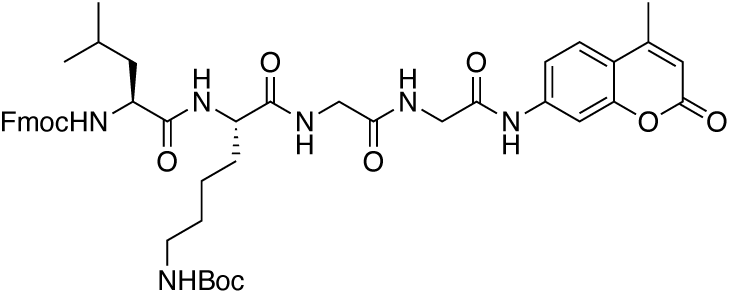

### Synthesis of S6

A solution of **S5** (22.7 mg, 26.6 *µ*mol, 1.0 equiv) in 10% Piperidine:CH_2_Cl_2_ (2 mL) was stirred for 15 min at 23 °C and followed with solvent removal by rotary evaporation. Dilution with DMF (5 mL) and solvent removal by rotary evaporation was repeated two more times to remove residual piperidine. The resultant crude was resuspended in DMF (2 mL), and *i*-Pr_2_NEt (18.5 *µ*L, 0.11 mmol, 4 equiv) and acetic anhydride (10.0 *µ*L, 0.11 mmol, 4 equiv) were added at 23 °C. The reaction mixture was stirred for 1 h at 23 °C, diluted by EtOAc (20 mL), and then washed by aqueous HCl (pH ∼2-3). The aqueous layer was further extracted by EtOAc (2 × 20 mL), the combined organic layers were dried over Na_2_SO_4_, and then evaporated to give a crude product. Purification by column chromatography (silica gel, 2-15% MeOH in 1:1 EtOAc:CH_2_Cl_2_) yielded **S6** (9.7 mg, 56% yield). **S6**: ^1^H NMR (400 MHz, DMSO-*d*_6_) δ 10.18 (s, 1 H), 8.39–8.30 (m, 1 H), 8.26 (t, *J* = 5.8 Hz, 1 H), 8.06 (d, *J* = 7.0 Hz, 1 H), 7.98 (d, *J* = 8.1 Hz, 1 H), 7.80 (d, *J* = 2.1 Hz, 1 H), 7.73 (d, *J* = 8.7 Hz, 1 H), 7.54 (dd, *J* = 8.7, 2.1 Hz, 1 H), 6.79–6.68 (m, 1 H), 6.27 (d, *J* = 1.3 Hz, 1 H), 4.29 (q, *J* = 7.6 Hz, 1 H), 4.16 (q, *J* = 6.5 Hz, 1 H), 3.94 (s, 2 H), 3.84–3.67 (m, 2 H), 2.86 (t, *J* = 6.4 Hz, 2 H), 2.40 (d, *J* = 1.3 Hz, 3 H), 1.83 (s, 3 H), 1.73–1.45 (m, 4 H), 1.36 (s, 11 H), 1.23 (s, 3 H), 0.78 (dd, *J* = 6.6, 3.9 Hz, 6 H); HRMS (ESI) calculated for C_33_H_48_N_6_O_9_ [M^+^] 672.3483, found 672.3502.

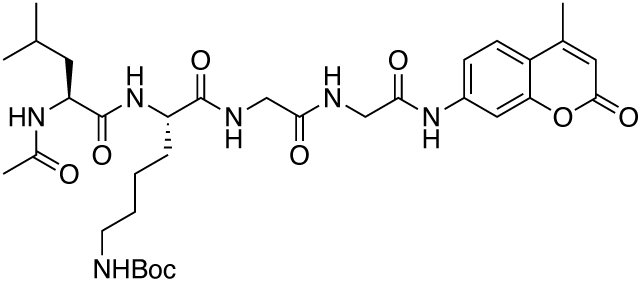

### Synthesis of CV-2

A solution of **S6** (11.2 mg, 16.6 *µ*mol, 1.0 equiv) in 20% TFA:CH_2_Cl_2_ (2 mL) was stirred at 23 °C for 30 min, followed by dilution with CH_2_Cl_2_ (5 mL) and solvent removal by rotary evaporation. Dilution with CH_2_Cl_2_ (5 mL) and solvent removal by rotary evaporation was repeated two more times to remove residual TFA. The crude was dissolved in DMSO to generate 20 mM stock of **CV-2**. Purity was assayed by LC/MS. **CV-2**: HRMS (ESI) calculated for C_28_H_40_N_6_O_7_ [M^+^] 572.2958, found 572.2956.

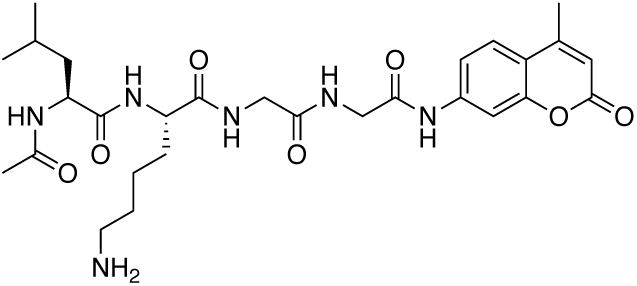

### Cells and Virus

Vero E6 cells (ATCC) were infected under biosafety level 3 conditions with SARS-CoV-2 (nCoV/Washington/1/2020, kindly provided by the National Biocontainment Laboratory, Galveston, TX).

### Immunostaining against Spike protein

Immunostaining was performed on 10% NBF fixed SARS-CoV-2 infected cells in 96-well plate. After fixation, 10% NBF was removed, and cells were washed with PBS, followed by washing with PBS-T (0.1% Tween 20 in PBS), and then blocked for 30 min with PBS containing 1% BSA at 23 °C. After blocking, endogenous peroxidases were quenched by 3% hydrogen peroxide for 5 min. Then, cells were washed with PBS and PBS-T and incubated with a monoclonal mouse-anti-SARS-CoV-2 spike antibody (GeneTex, 1:1000) in PBS containing 1% BSA overnight at 4 °C. Primary antibody was washed with PBS and PBS-T and then cells were incubated in secondary antibody (ImmPRESS Horse Anti-Mouse IgG Polymer Reagent, Peroxidase; Vector Laboratories) for 60 min at 23 °C. After washing with PBS for 10 min, color development was achieved by applying diaminobenzidine tetrahydrochloride (DAB) solution (Metal Enhanced DAB Substrate Kit; ThermoFisher Scientific) for 30 min and observed by light microscopy for the percentage of cells that were Spike positive.

## Supplemental Figure Legends

**Supplemental Figure 1.**
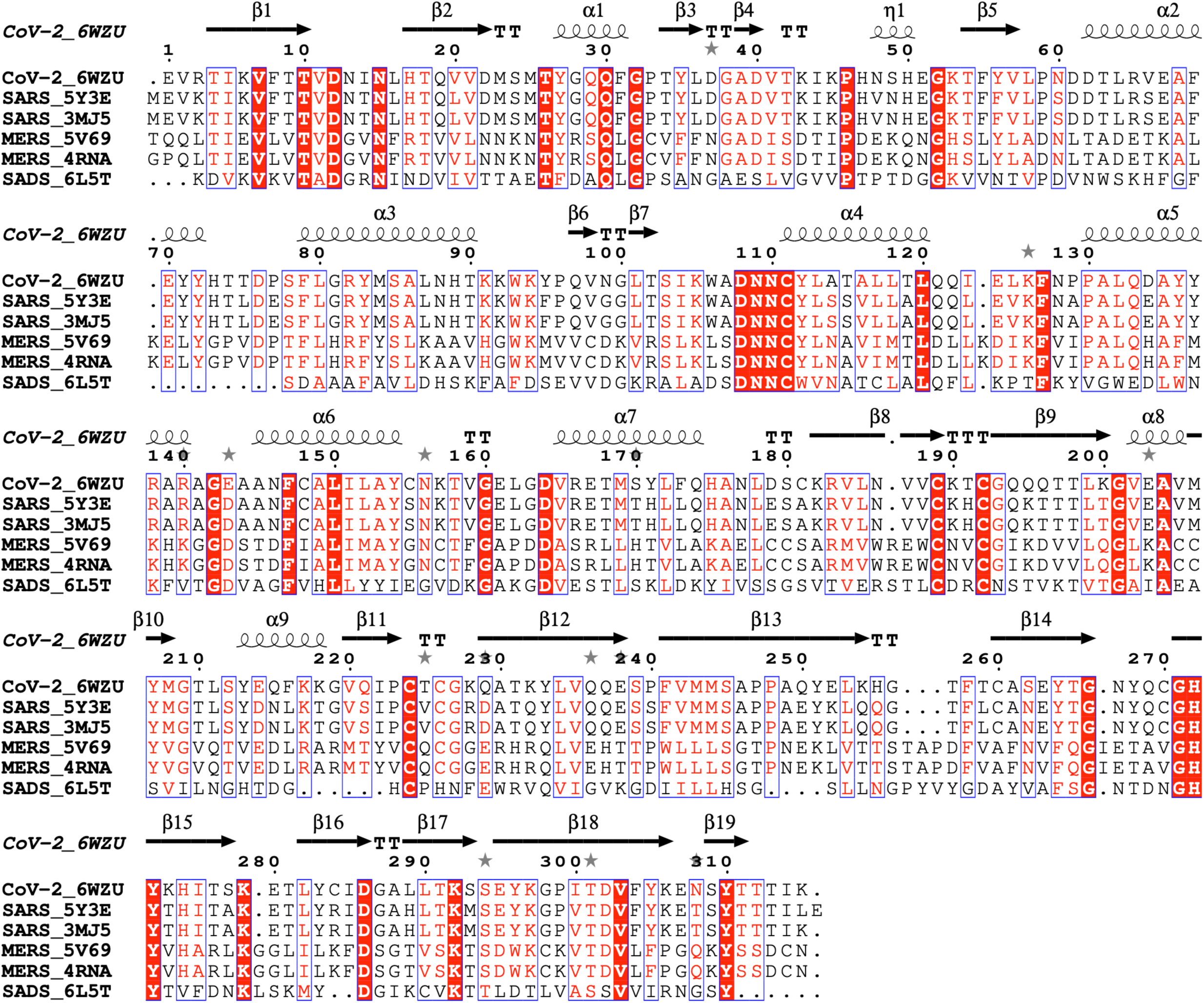
Sequence alignment of SARS-CoV-2 PLpro homologues from CoV-2, SARS, MERS and SADS coronaviruses with structures available in the PDB: SARS CoV-2 (PDB id: 6WZU), SARS CoV (PDB ids: 5Y3E and 3MJ5), MERS CoV (PDB id: 5V69 and 4RNA) and SADS CoV (PDB id: 6L5T). The secondary-structure elements are labelled for SARS-CoV-2 PLpro.

**Supplemental Figure 2A.**
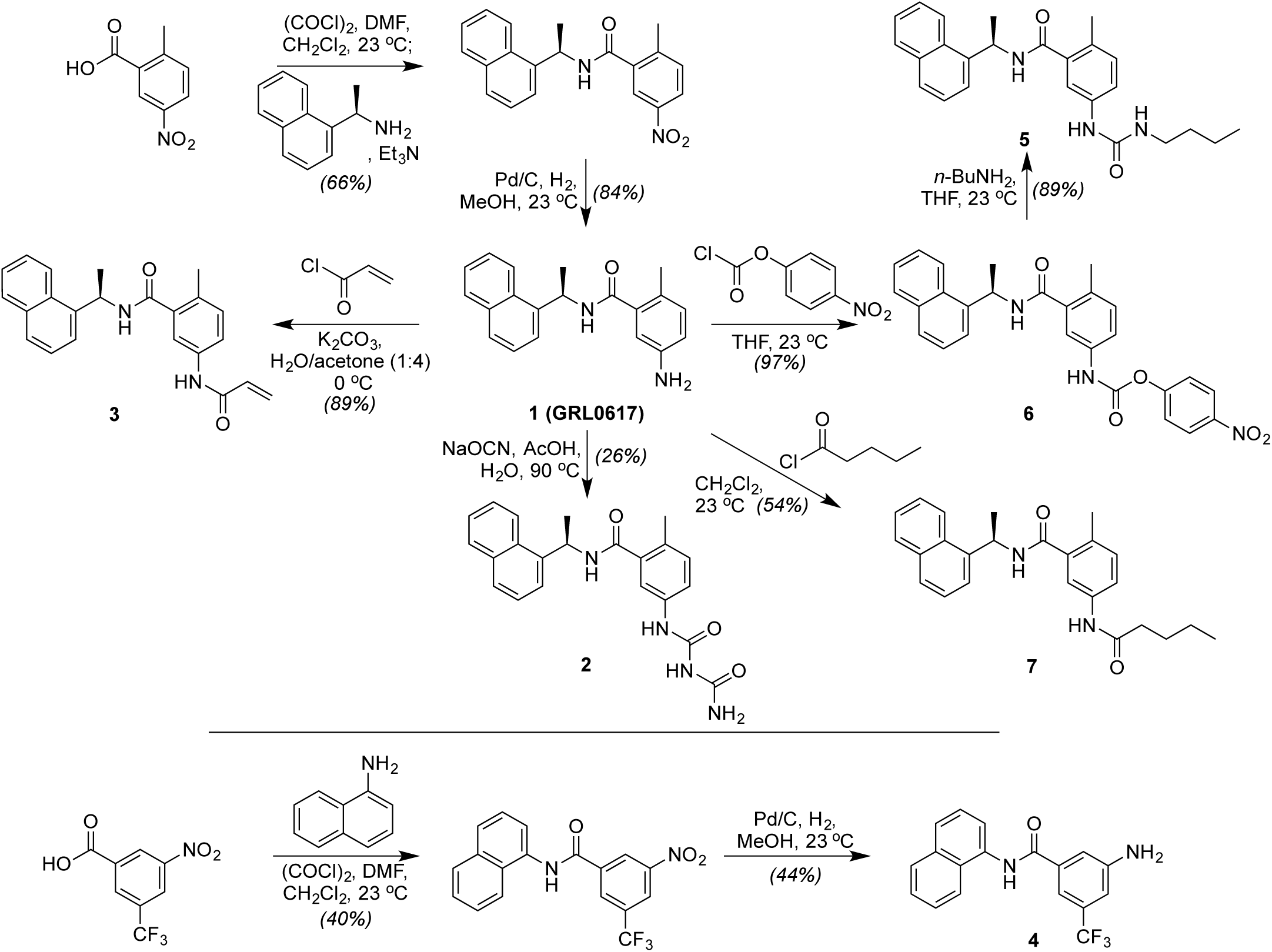
Synthesis of inhibitors 1-7 starting from commercially available materials.

**Supplemental Figure 2B.**
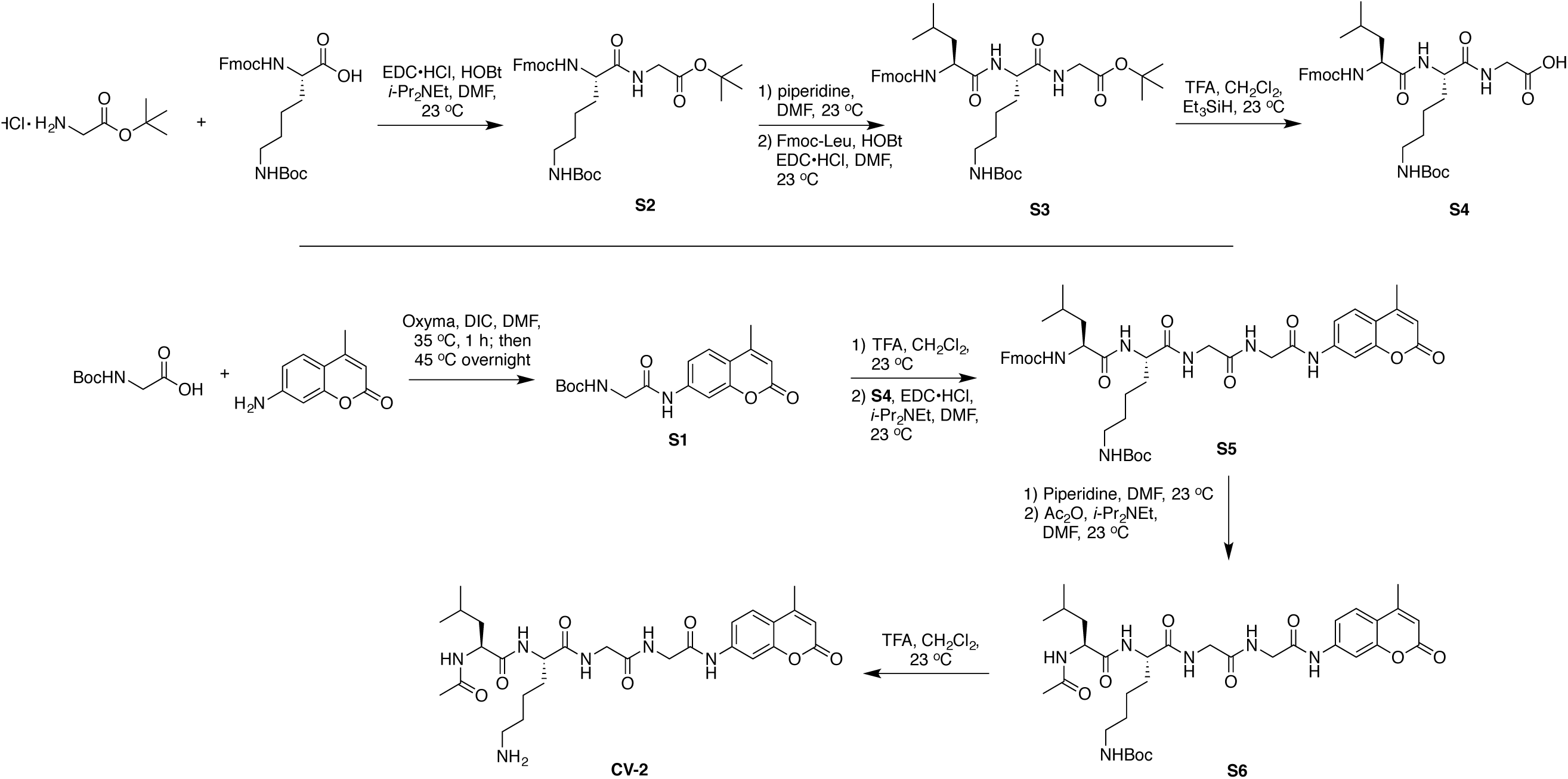
Synthesis of CV-2.

**Supplemental Figure 3:**
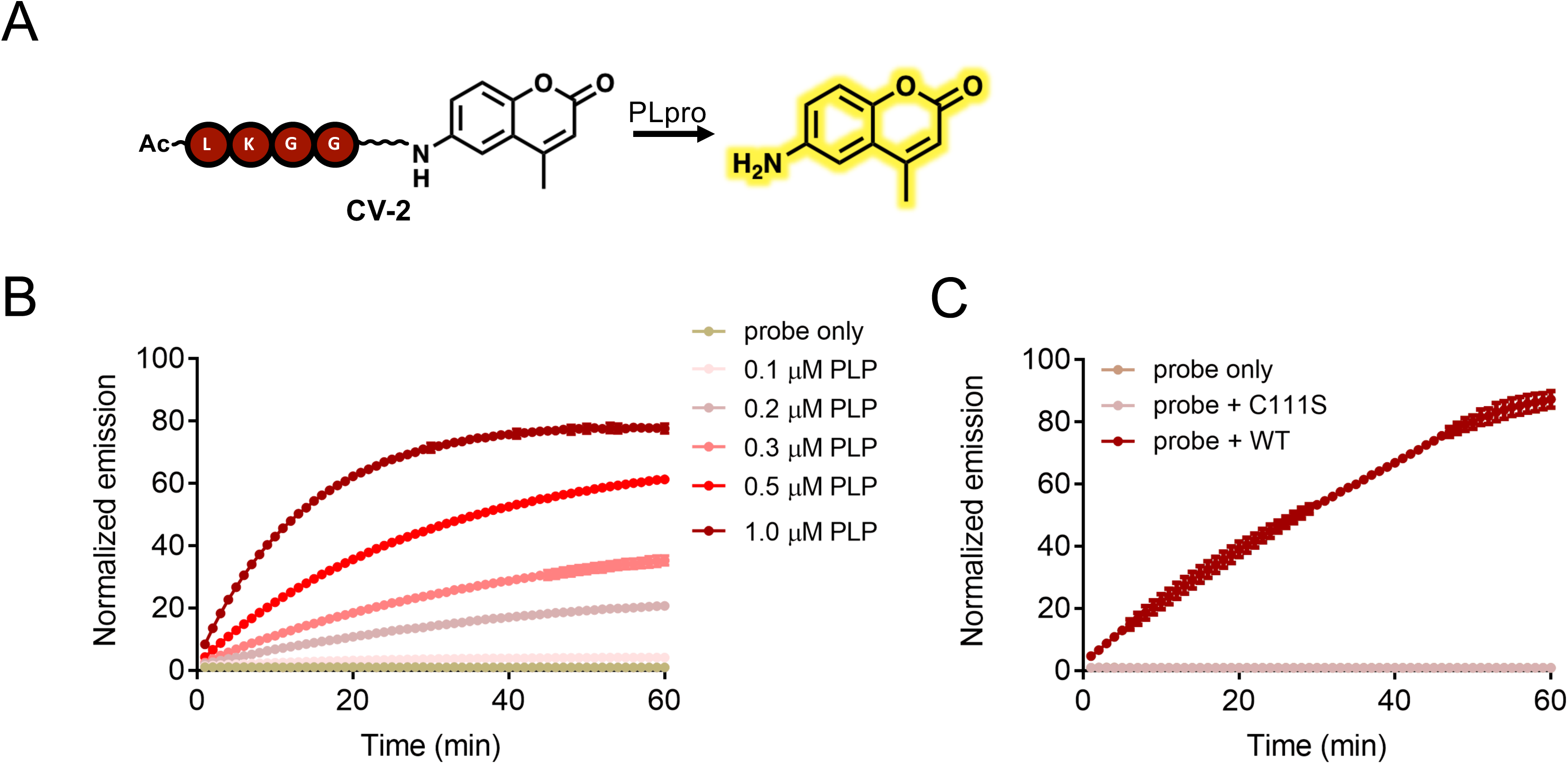
Biochemical assay for PLpro. A) CV-2 features a PLpro peptide substrate tethered to a profluorescent molecule which is cleaved when enzymatic activity of PLpro releases a fluorescent product. B) CV-2 (40 *µ*M) incubated with varying amounts of PLpro and fluorescence quantified over time by plate reader. C) Comparison of wild-type to active site mutant (C111S) shows biochemical assays reports on active proteolysis of PLpro.

**Supplementary Figure 4.**
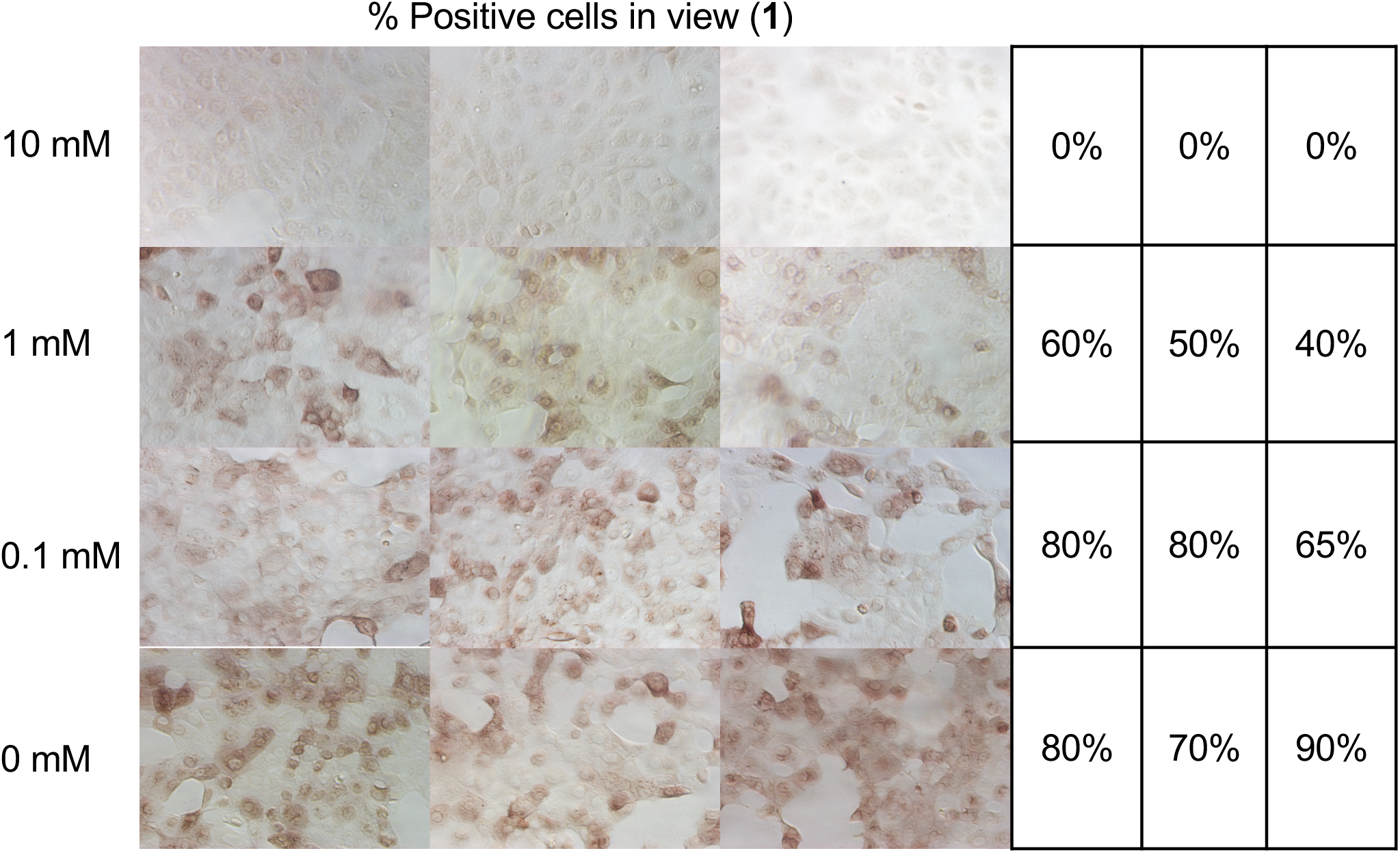
Whole cell assay for compound **1.**

**Supplemental Figure 5.**
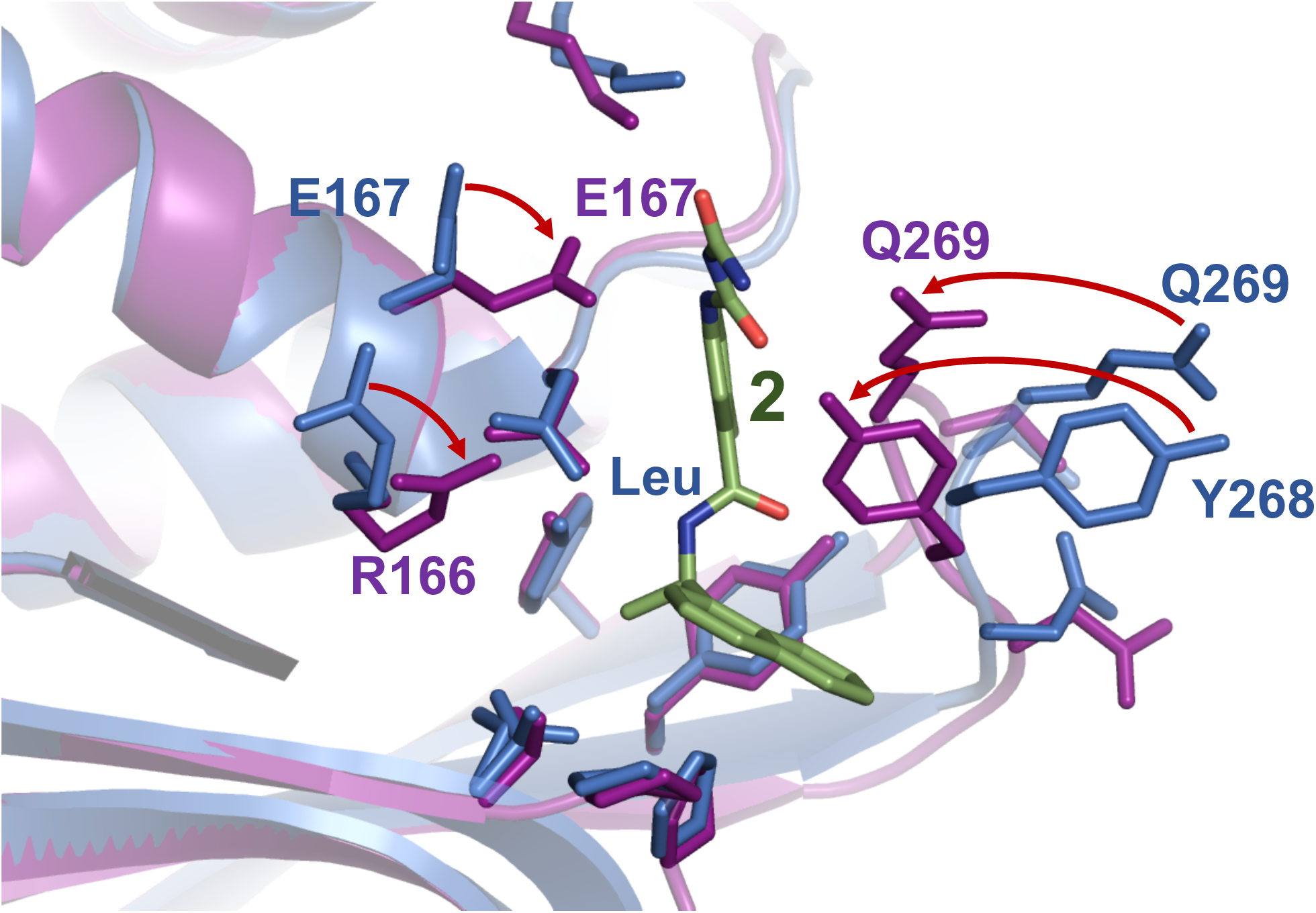
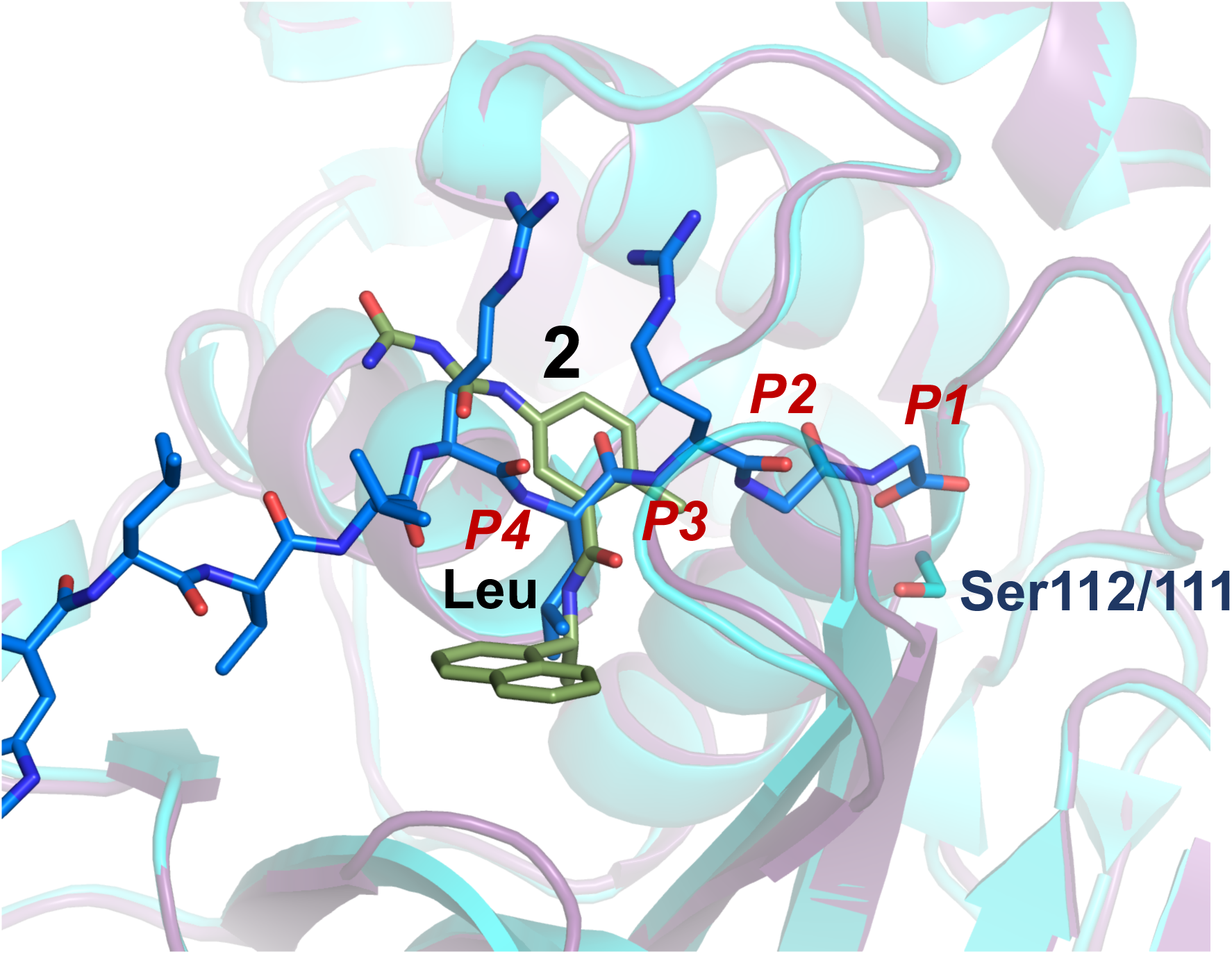
PLpro ligand binding. A) Superposition of PLpro ligand-binding sites of the unliganded WT protein structure (shown in blue, 6WZU id: PDB) and the structure with compound **2** (in magenta with the ligand in green, PDB id: 7JIT). B) Structure superposition of ligand-binding sites of PLpro compound **2** complex (in magenta with ligand in green, PDB id: 7JIT) and SARS-CoV PLpro C112S mutant in complex with ubiquitin (shown in blue with C-terminal part of ubiquitin shown as blue-navy blue sticks, PDB id: 4M0W).

---

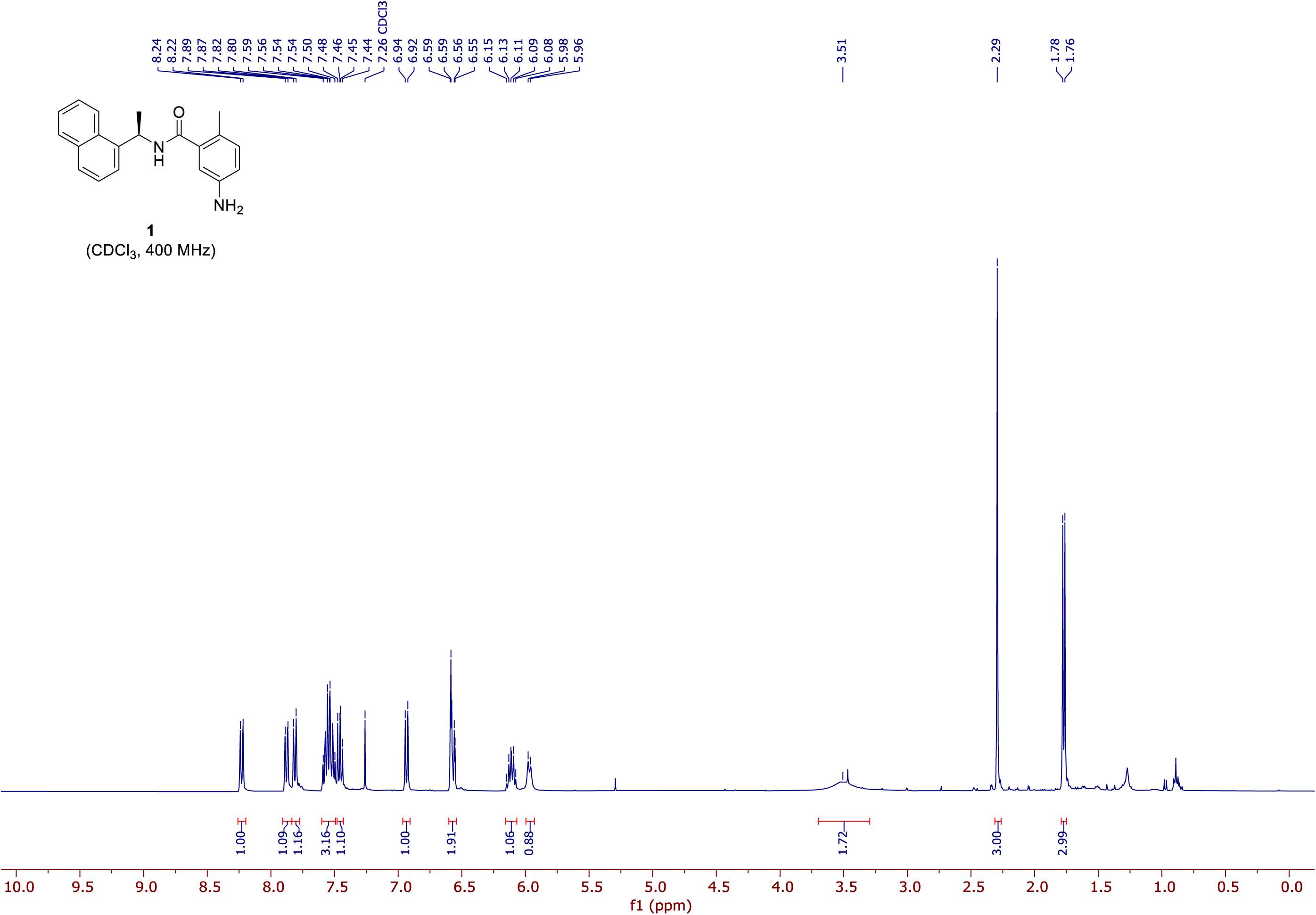

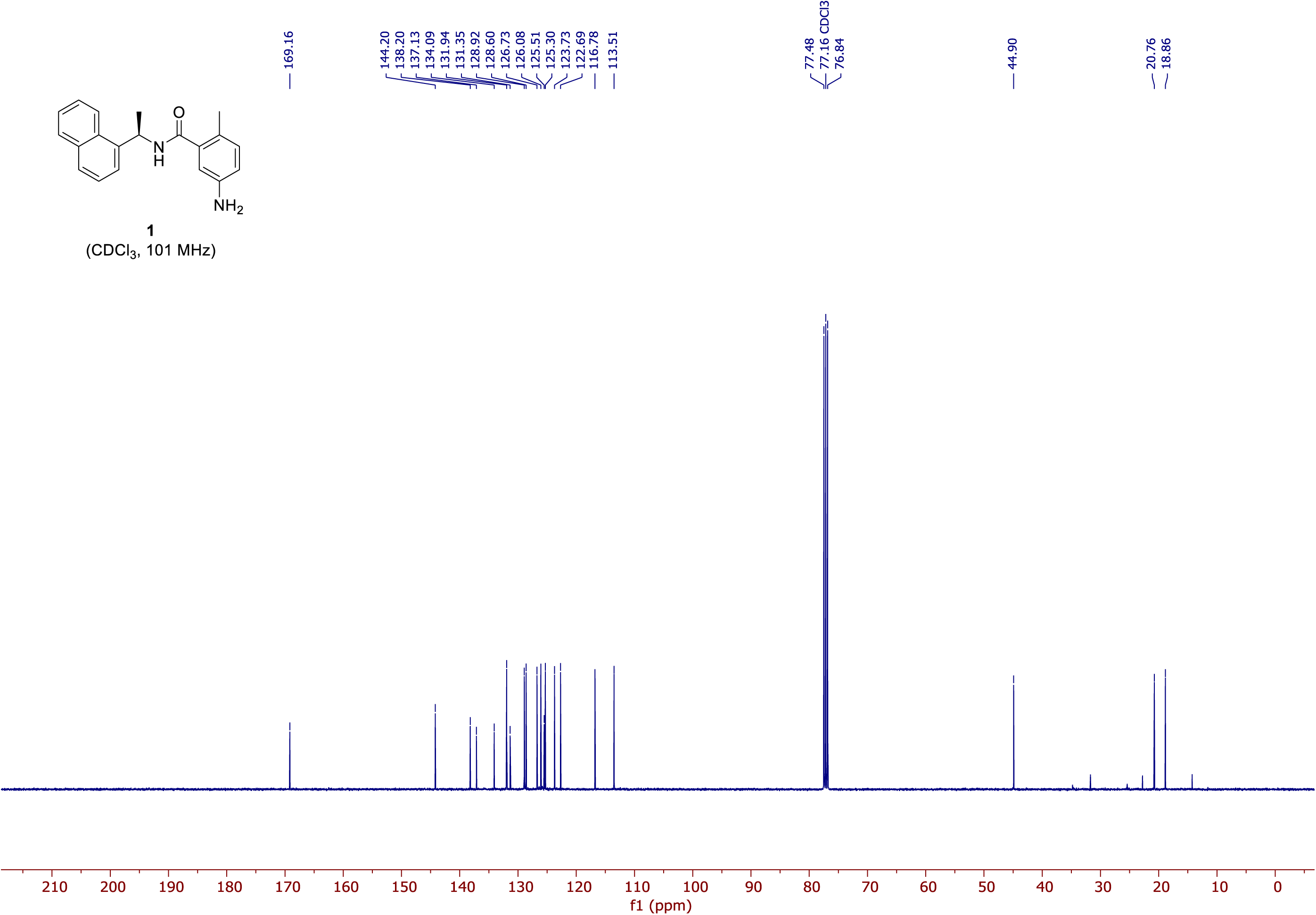

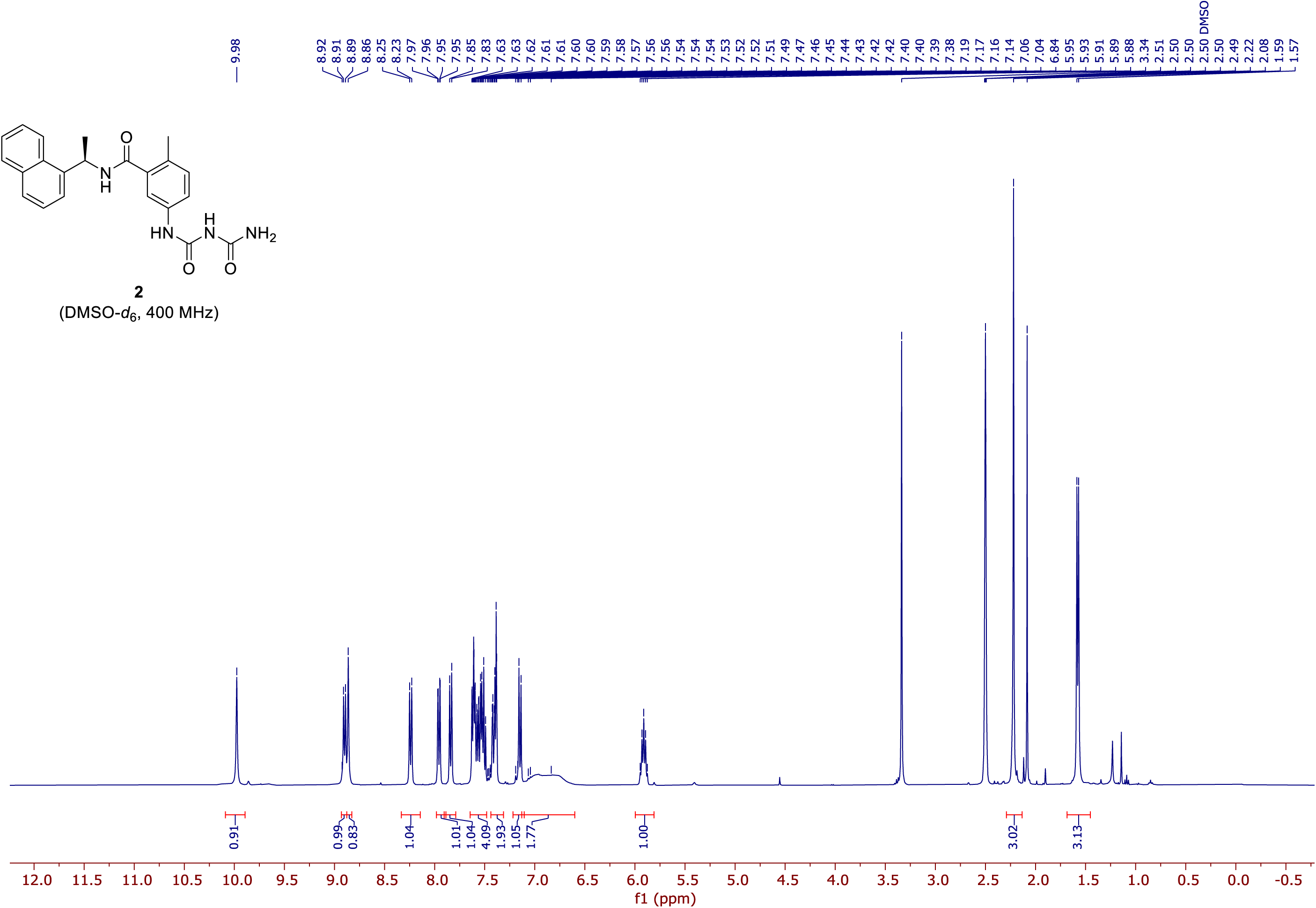

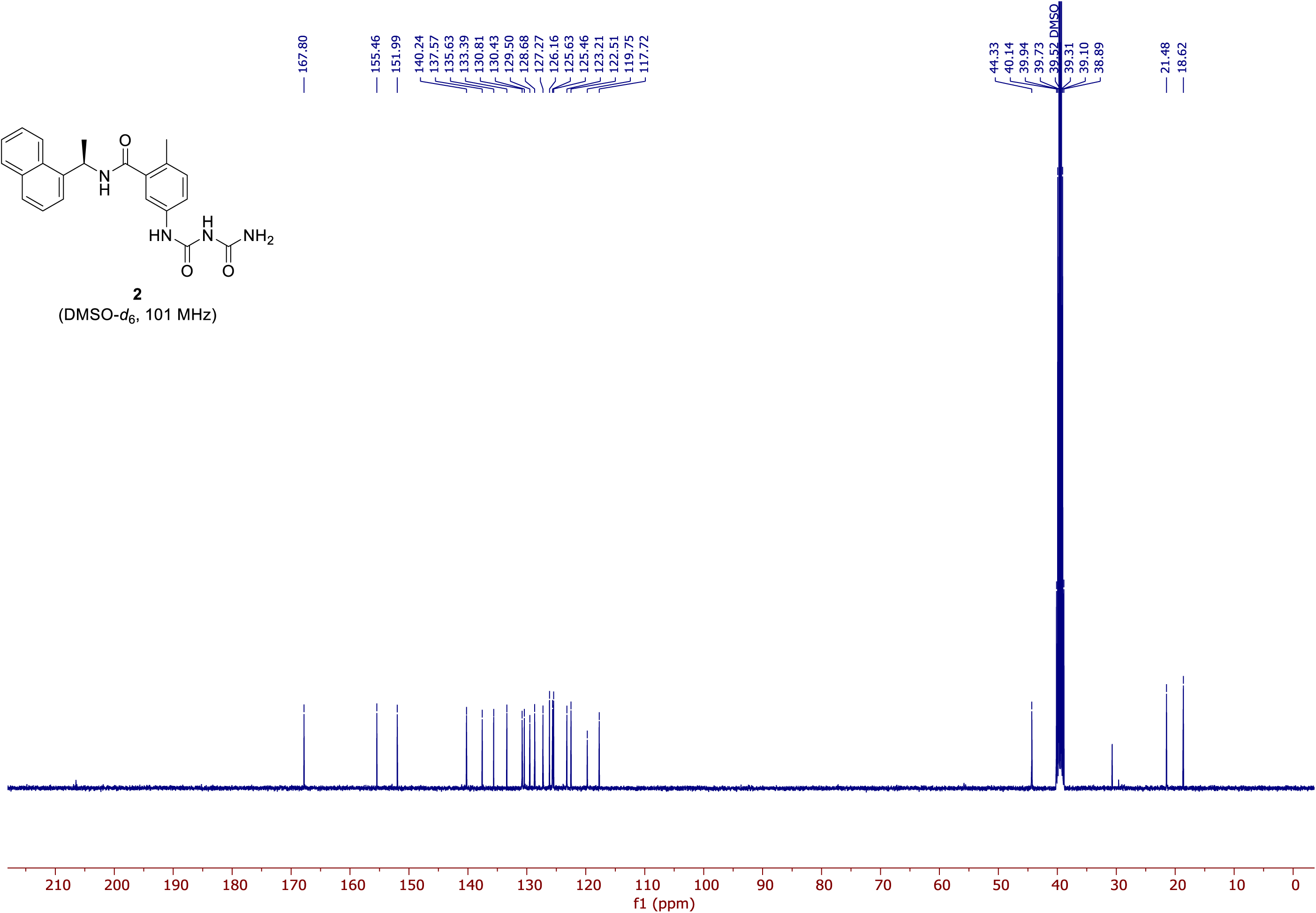

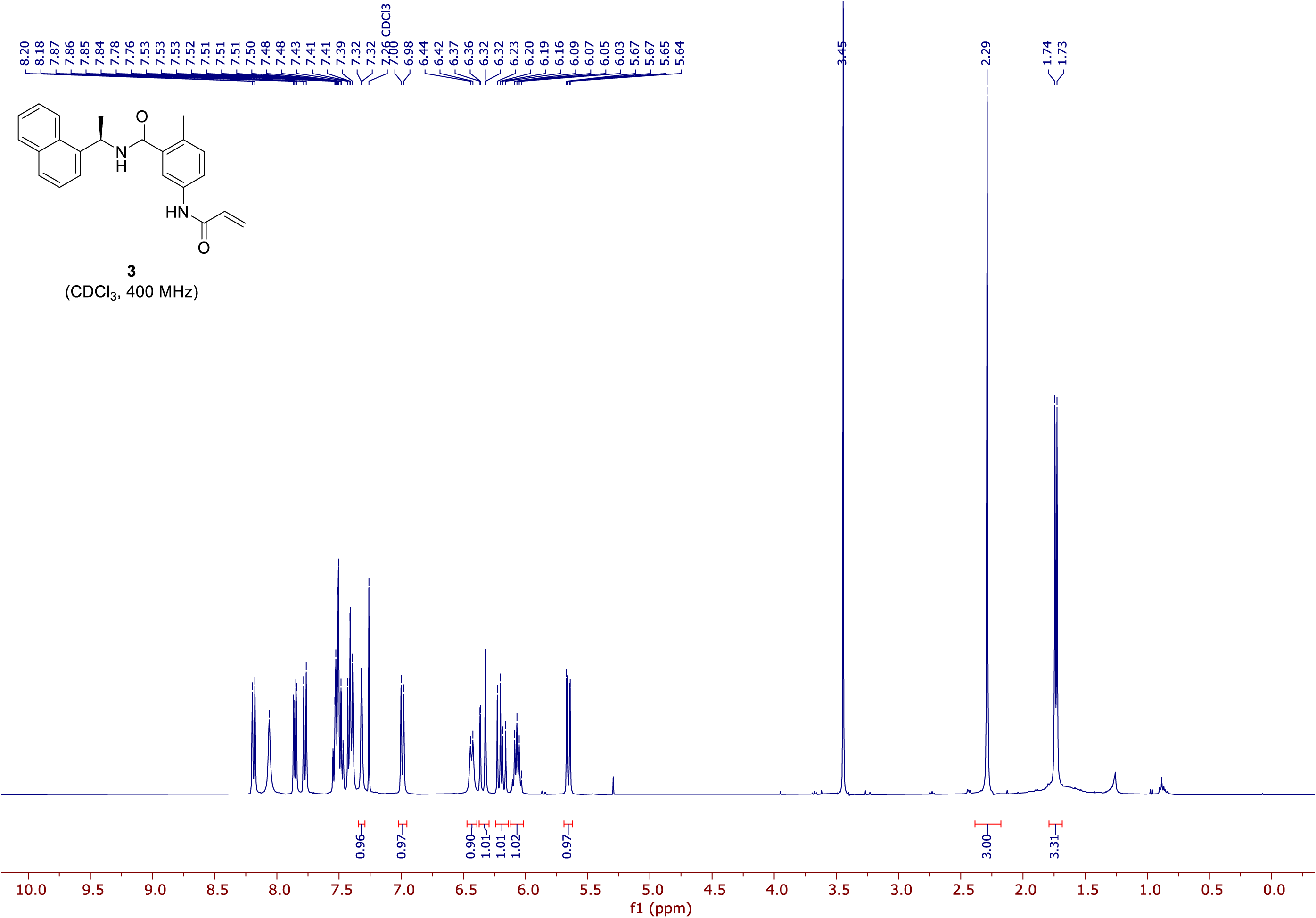

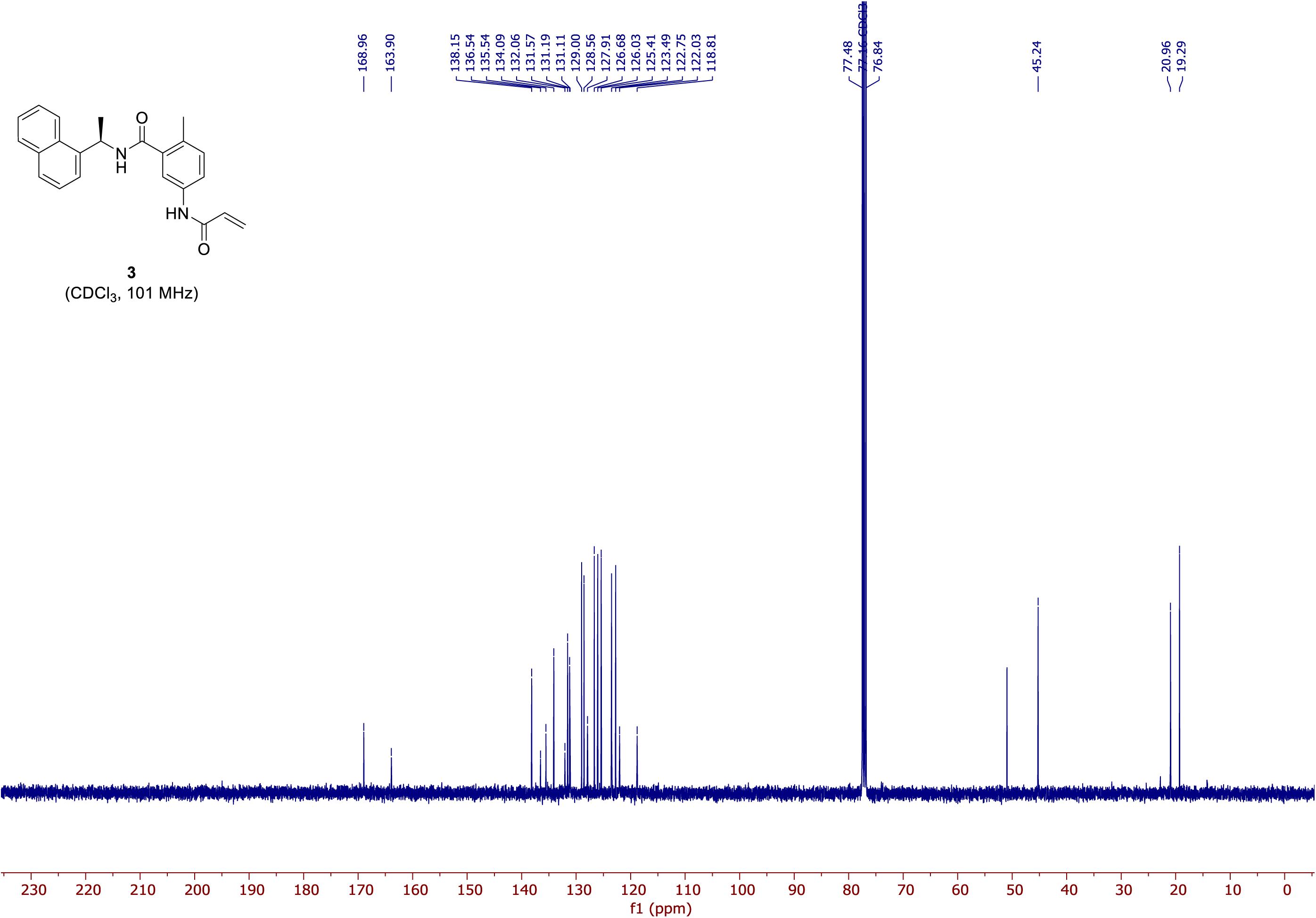

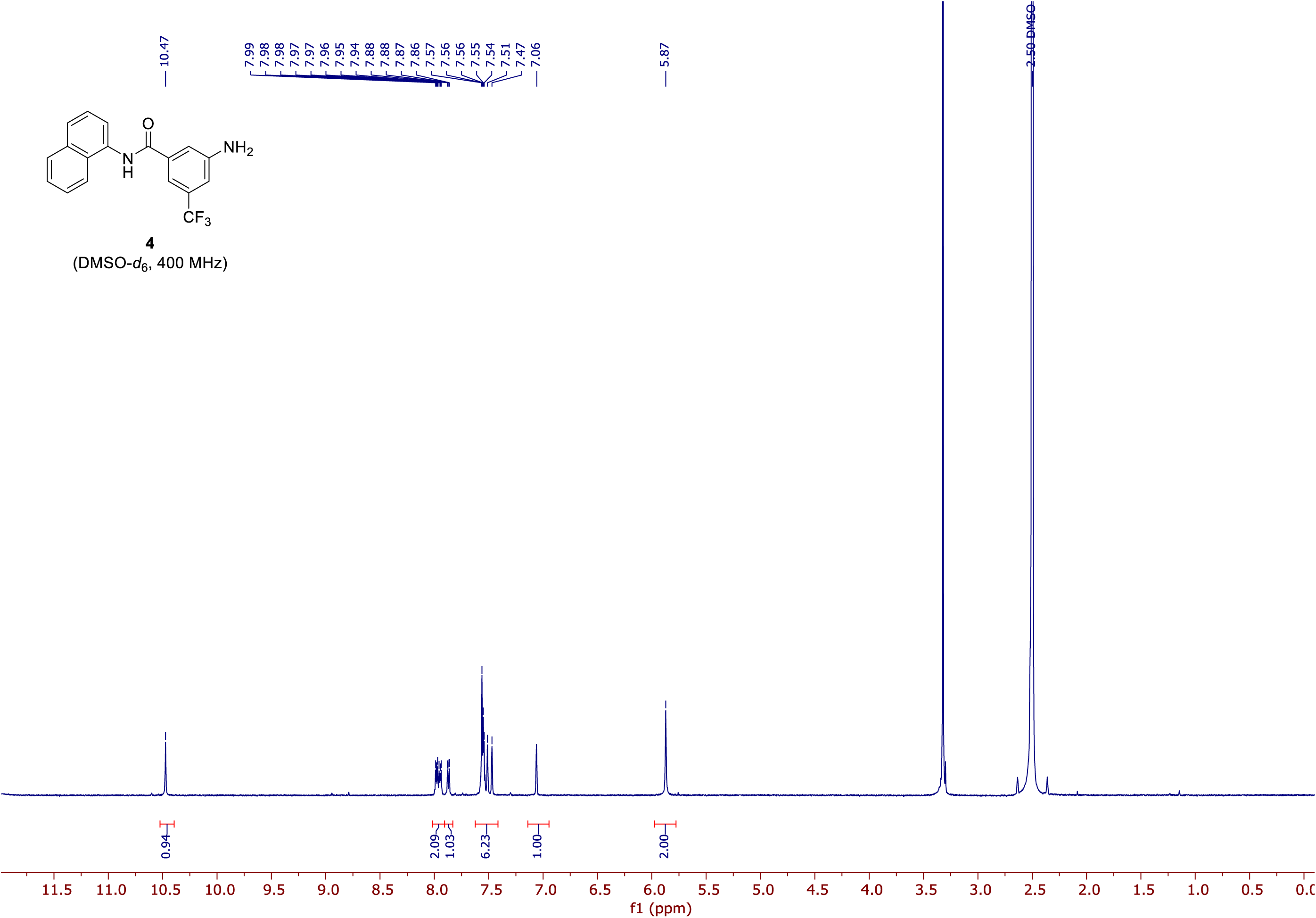

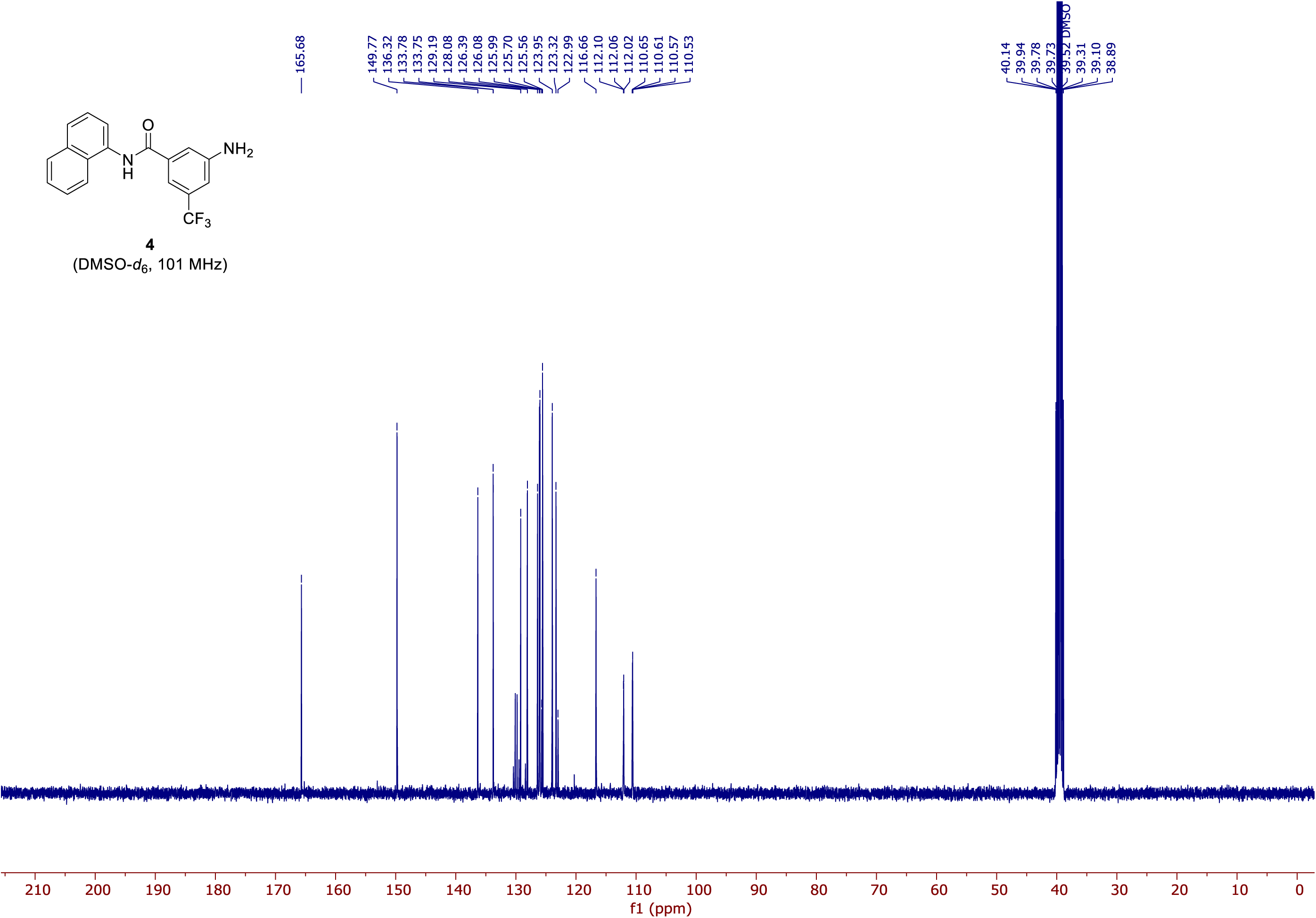

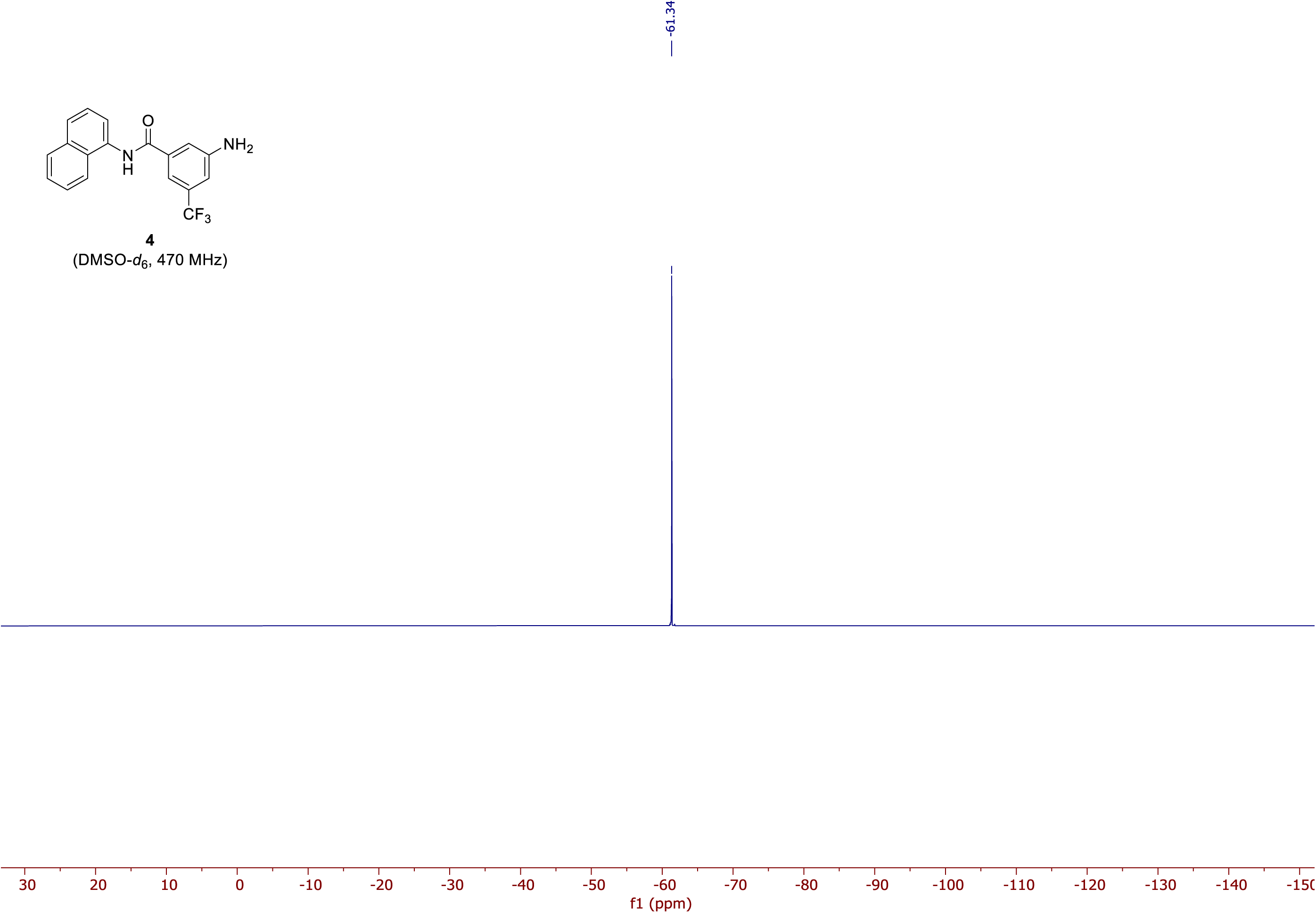

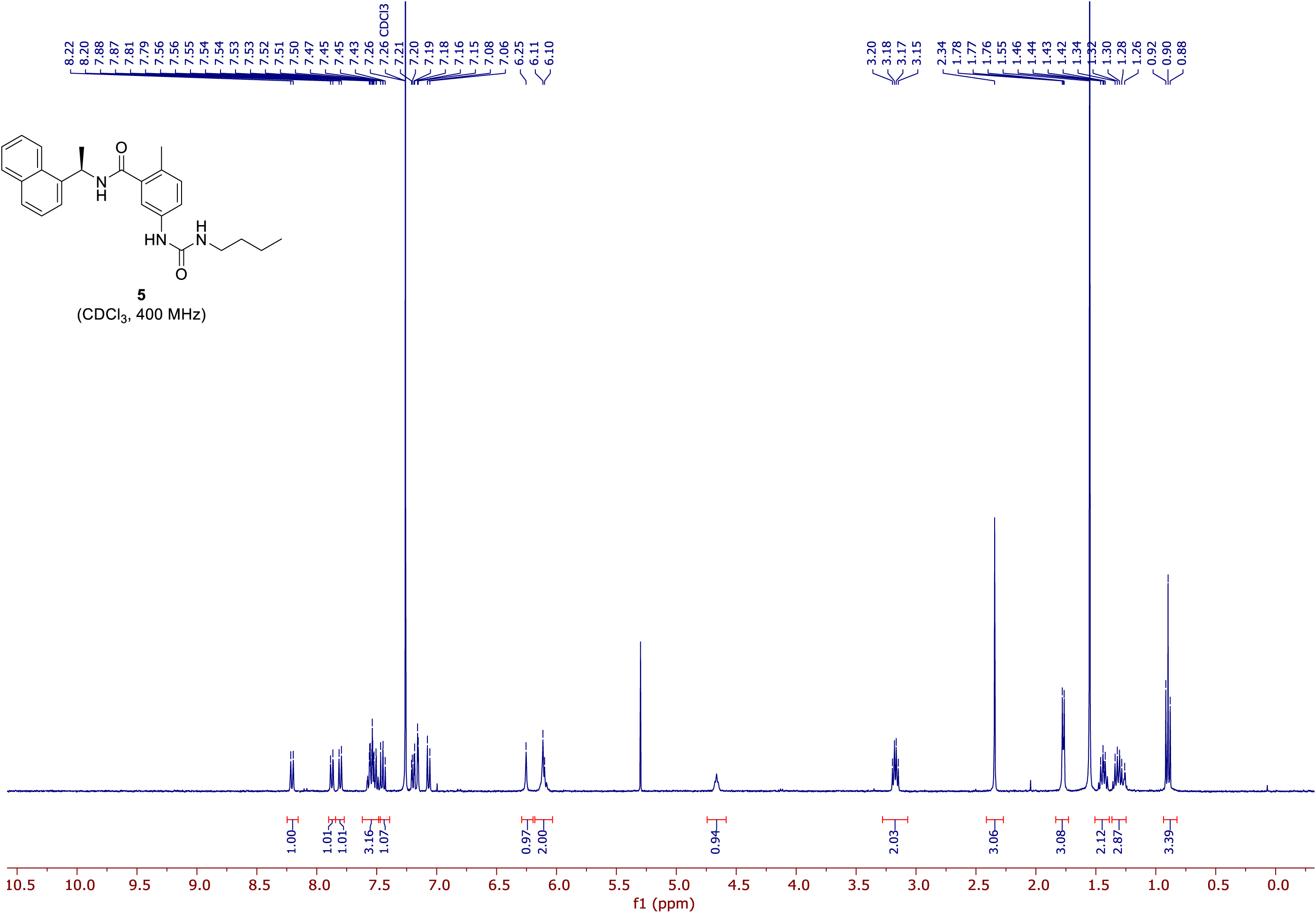

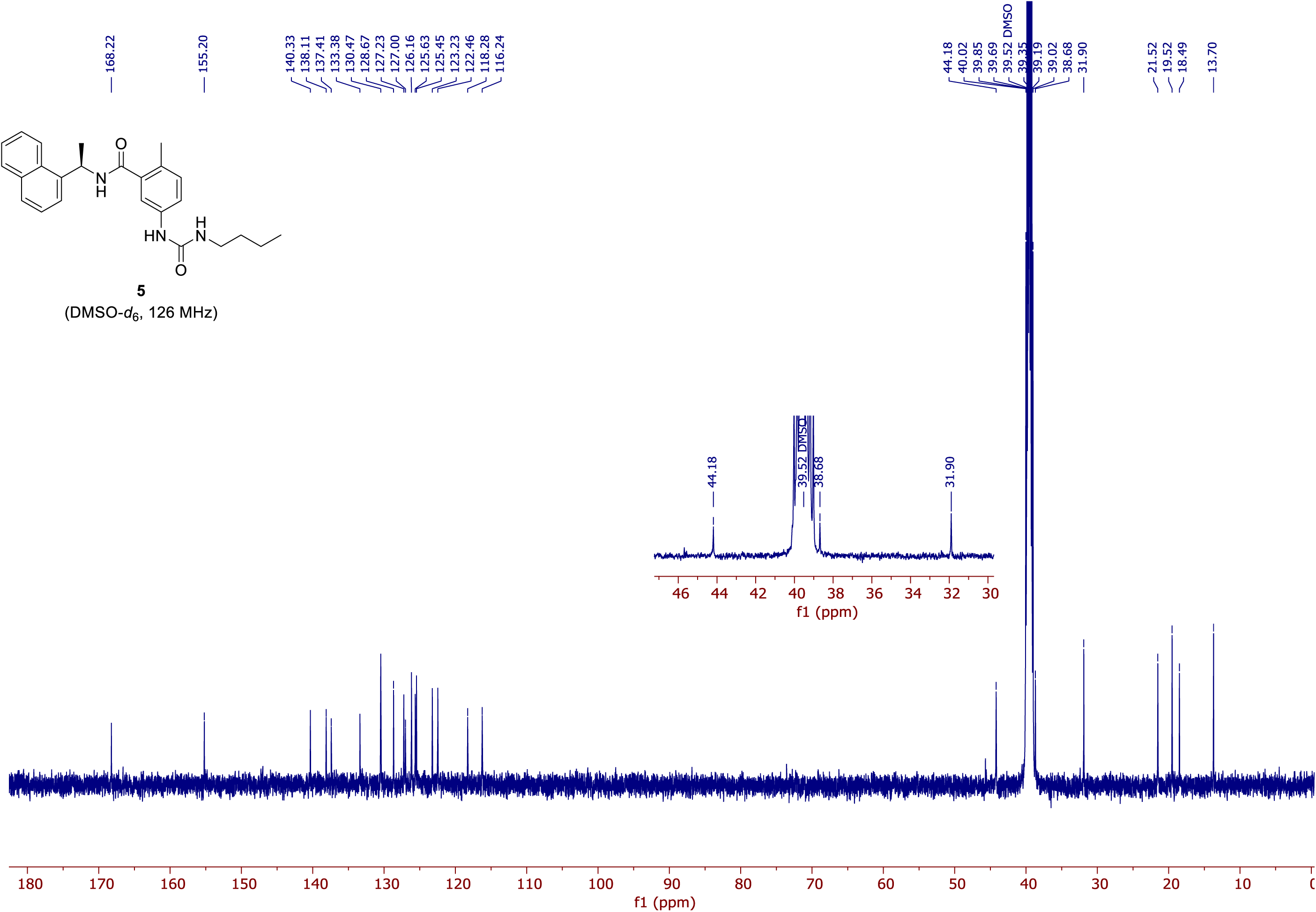

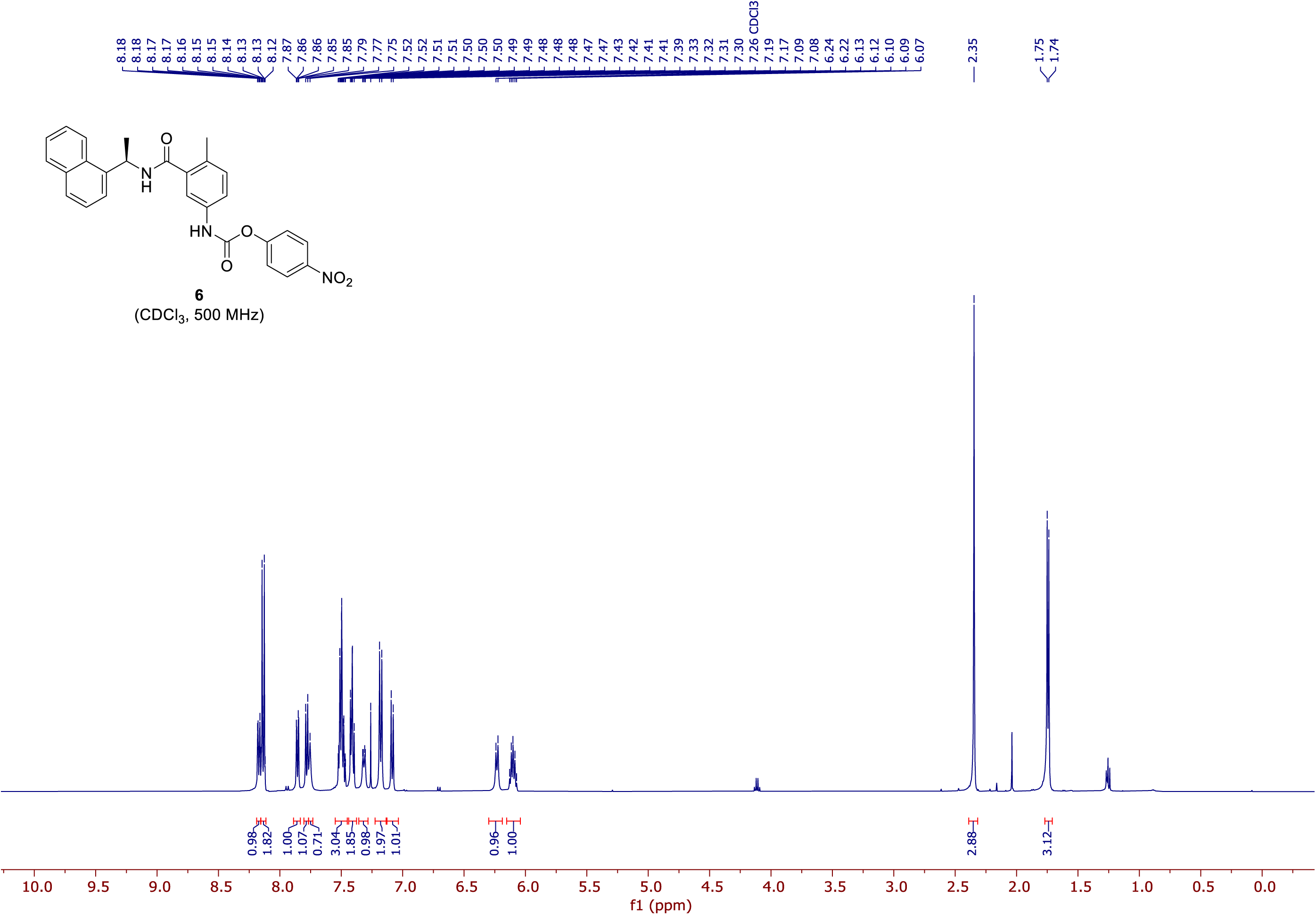

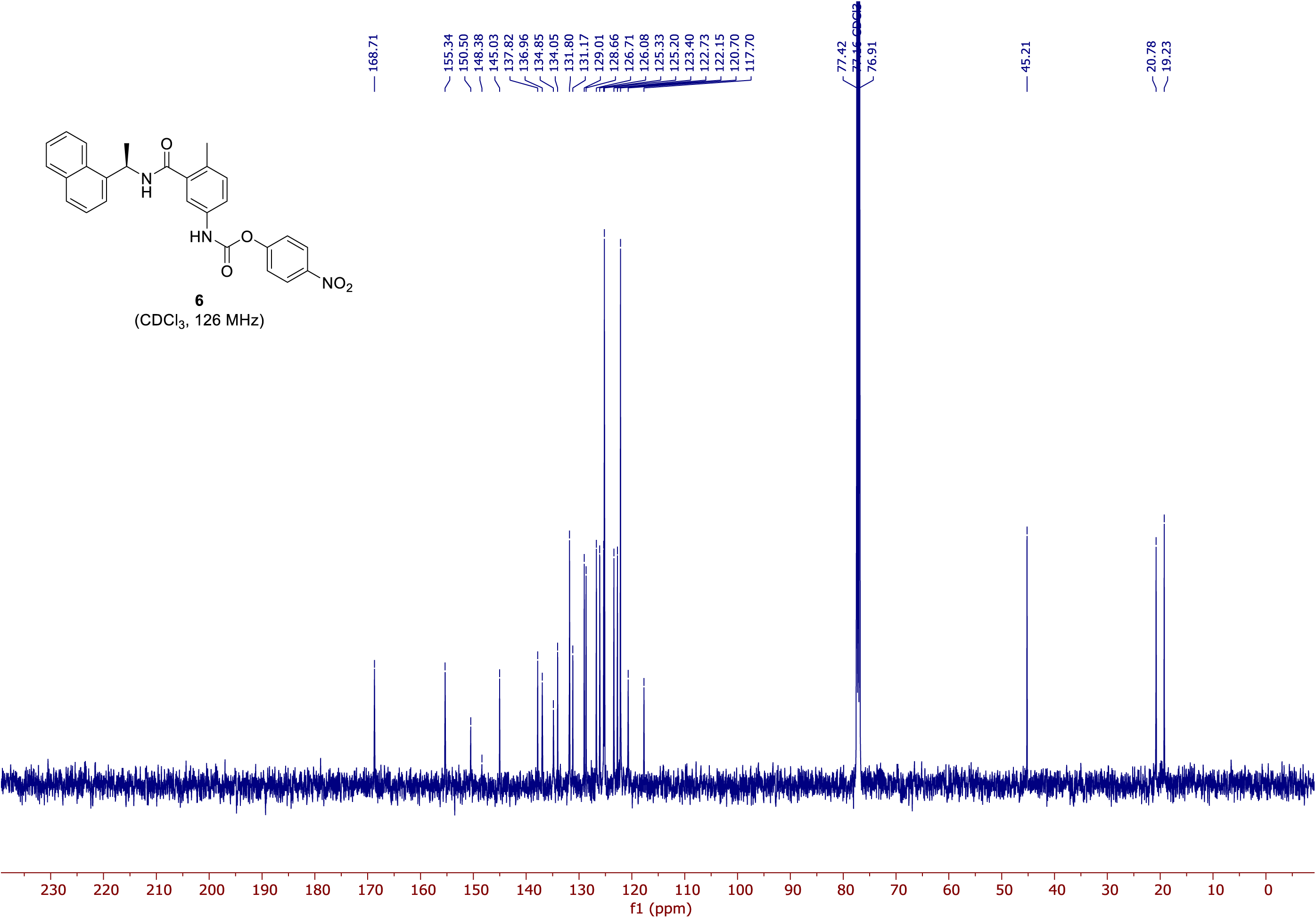

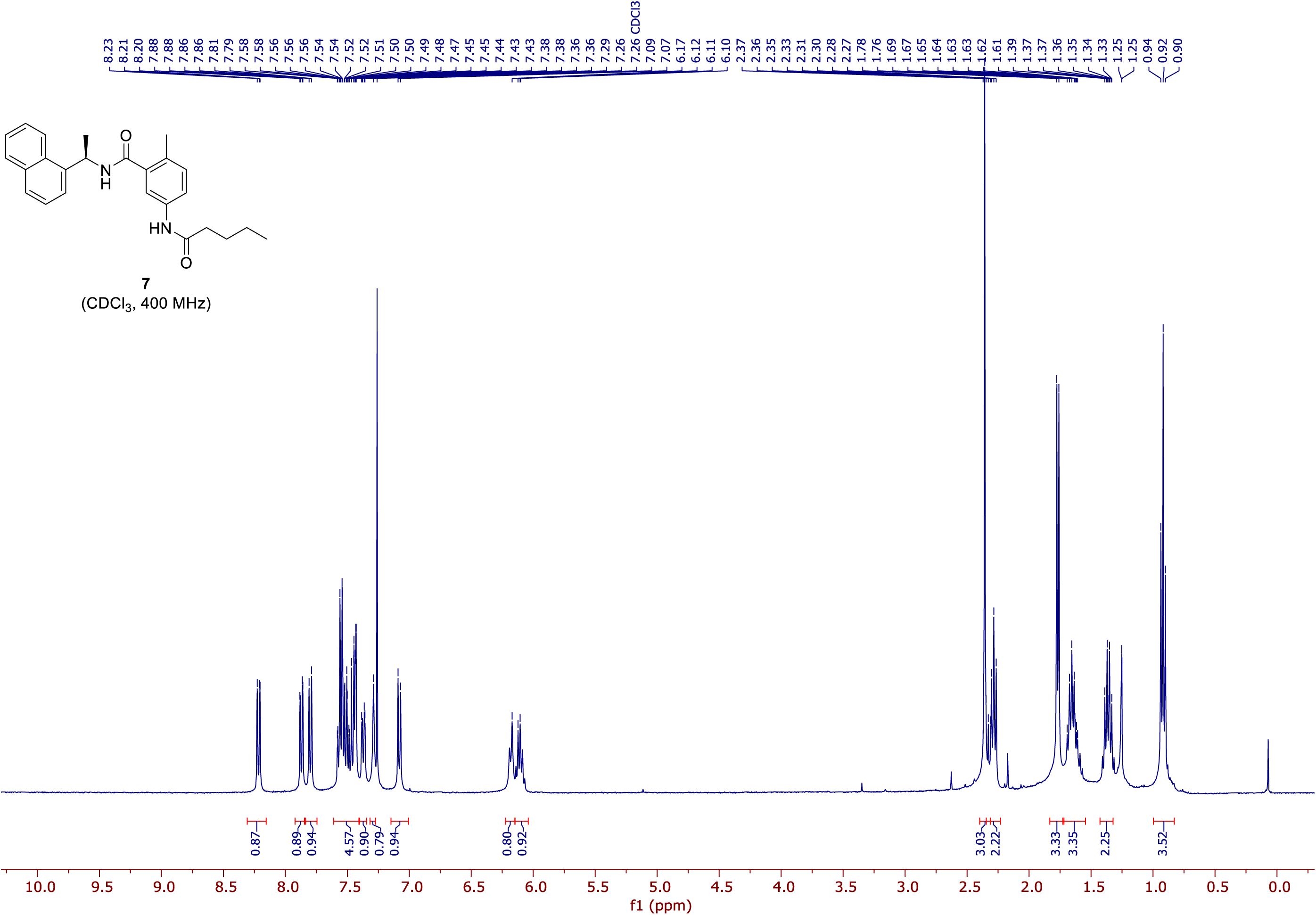

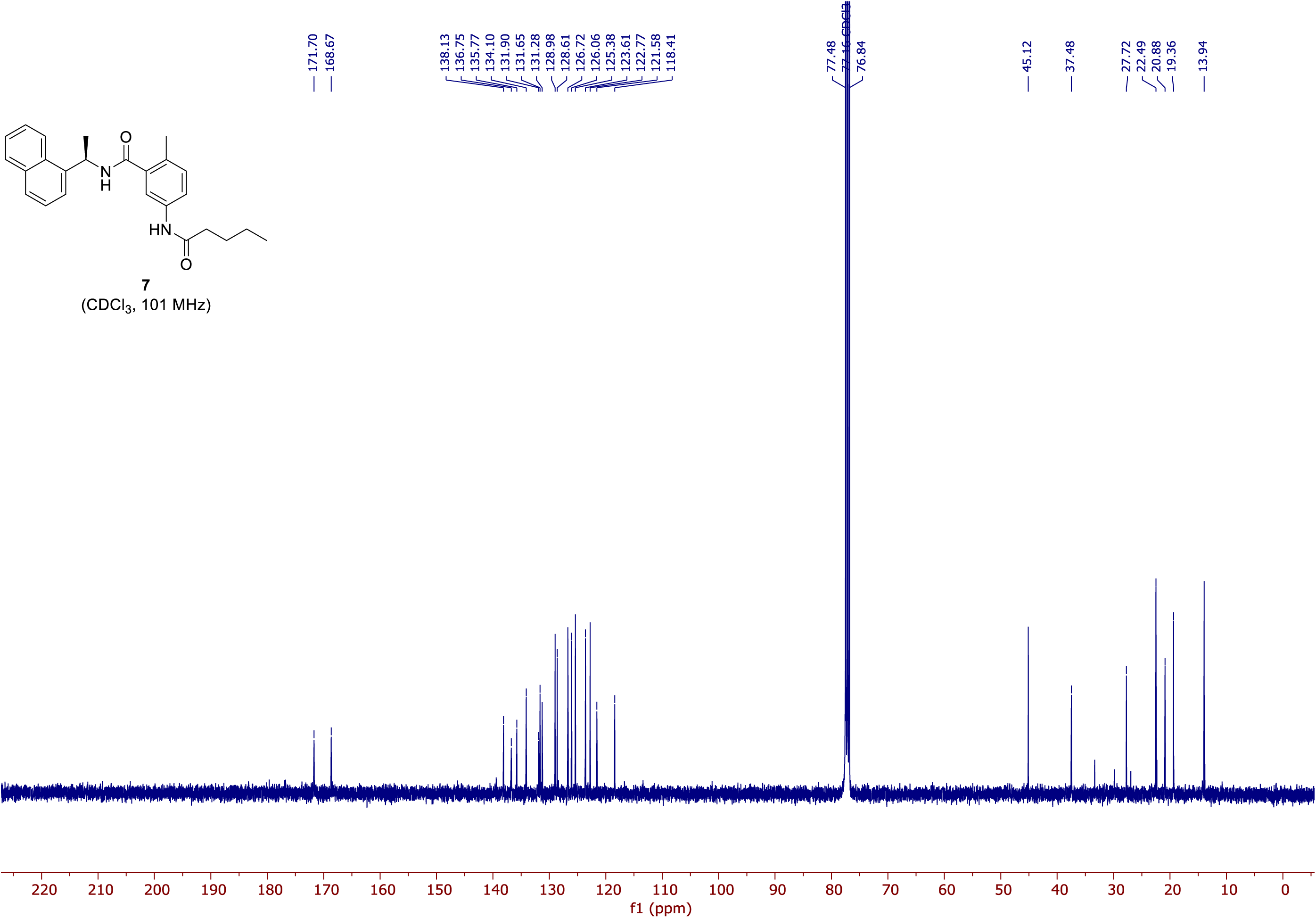

